# *Tmod2* is a regulator of cocaine responses through control of striatal and cortical excitability, and drug-induced plasticity

**DOI:** 10.1101/648295

**Authors:** Arojit Mitra, Sean P. Deats, Price E. Dickson, Jiuhe Zhu, Justin Gardin, Brian J. Nieman, R. Mark Henkelman, Nien-Pei Tsai, Elissa J. Chesler, Zhong-Wei Zhang, Vivek Kumar

## Abstract

Drugs of abuse induce neuroadaptations, including synaptic plasticity, that are critical for transition to addiction, and genes and pathways that regulate these neuroadaptations are potential therapeutic targets. *Tropomodulin 2* (*Tmod2*) is an actin-regulating gene that plays an important role in synapse maturation and dendritic arborization and has been implicated in substance-abuse and intellectual disability in humans. Here we mine the KOMP2 data and find that *Tmod*2 knockout mice show emotionality phenotypes that are predictive of addiction vulnerability. Detailed addiction phenotyping showed that *Tmod2* deletion does not affect the acute locomotor response to cocaine administration. However, sensitized locomotor responses are highly attenuated in these knockouts, indicating perturbed drug-induced plasticity. In addition, *Tmod2* mutant animals do not self-administer cocaine indicating lack of hedonic responses to cocaine. Whole brain MR imaging shows differences in brain volume across multiple regions although transcriptomic experiments did not reveal perturbations in gene co-expression networks. Detailed electrophysiological characterization of *Tmod2* KO neurons, showed increased spontaneous firing rate of early postnatal and adult cortical and striatal neurons. Cocaine-induced synaptic plasticity that is critical for sensitization is either missing or reciprocal in *Tmod2* KO nucleus accumbens shell medium spiny neurons, providing a mechanistic explanation of the cocaine response phenotypes. Combined, these data provide compelling evidence that *Tmod2* is a major regulator of plasticity in the mesolimbic system and regulates the reinforcing and addictive properties of cocaine.

**Significance statement:** We identify, characterize, and establish tropomodulin 2 *(Tmod2),* an actin-regulating gene exclusively expressed in neurons, as an important regulator of addiction-related phenotypes. We show that *Tmod2*, knockout mice (*Tmod2 KO*) exhibit phenotypes that are predictive of addiction. In detailed addiction phenotyping, we find the *Tmod2* regulates cocaine sensitization and self-administration. We explore anatomical, transcriptional, electrophysiological mechanisms of this regulation. Combined these studies provide compelling evidence that *Tmod2* is critical for synaptic plasticity necessary for transition to addiction.

## Introduction

Multiple lines of evidence suggest that enduring functional and structural plasticity in reward-related brain regions are critical for transition to addiction (1, 2). Although the entire brain reward circuit is vulnerable to drug-induced functional and structural remodeling (3), it is relatively well characterized in nucleus accumbens. Functional plasticity broadly refers to changes in intrinsic and synaptic properties of existing synapses whereas structural plasticity includes morphological changes such as dendritic spine class and density alterations (4). Drug-induced functional plasticity in nucleus accumbens includes changes in intrinsic properties such as firing rate depression, or synaptic changes such as AMPAR-mediated long-term depression (5, 6). These plasticity events have been shown to facilitate initiation and maintenance of reward-related behaviors (5-8).

Along with functional plasticity, repeated drug administration is also thought to facilitate structural plasticity in the mesolimbic circuit. Repeated administration of psychostimulants correlates with increased spine density and dendritic complexity in the nucleus accumbens, ventral tegmental area, and prefrontal cortex neurons (8). Dendritic spines are actin-rich protrusions of the neural plasma membrane that are major postsynaptic sites of excitatory signaling in the brain and sub-cellular substrate of synaptic and structural plasticity. Perturbations in actin-assembly (9, 10), -capping (11), -disassociating (12), -nucleating (13) and -ADP/ATP exchange (14) proteins affects synaptic and structural plasticity. Previous studies have implicated actin dysregulation in morphine (15), cocaine (16), methamphetamine (17), alcohol (18) and nicotine (19) addiction. Generally, psychostimulant administration correlates with increases in spine numbers, spine complexity, and spine maturity (20). These structural changes occur through actin-mediated rearrangements (12). Consistent with these observations, non-muscle myosin IIB inhibitor, which mitigates spine maturation by inhibiting crosslinking and treadmilling of F-actin, has been shown effective in preventing relapse to methamphetamine (21). Our previous study demonstrated that a point mutation in actin-regulating cytoplasmic FMRP interacting protein-2 (*Cyfip2*) in mice decreases both acute and sensitized cocaine responses (13). However, a causal link between structural changes and behavioral effects of repeated exposure to drugs remains controversial (2). Genetic or pharmacological perturbations of factors that are known to regulate structural plasticity, such as kalirin-7, myocyte enhancer factor-2, and cyclin-dependent kinase-5, fail to attenuate behavioral locomotor sensitization in mice (22-24). Thus, the possibility has been raised that structural plasticity in transition to addiction is an “epiphenomenon” (2).

The neuron-specific, tropomyosin-dependent pointed end capping protein tropomodulin-2 (TMOD2), stabilizes actin filament by preventing depolymerization and elongation (25). Four isoforms of tropomodulin (*Tmod1-4*) are expressed with tissue specificity in mice. *Tmod1* and *Tmod3* are ubiquitously expressed, whereas *Tmod4* and *Tmod2* are restricted to muscle and neurons, respectively (26). Previous studies have established a role for TMOD2 in dendritic arborization (27) and spine maturation (28). During development, its expression coincides with generation, migration, and extension of cerebrum and cerebellum neurons during the prenatal period (29, 30). In addition to pointed end capping of actin filament, an *in vitro* protein interaction study shows actin-monomer binding and strong actin-nucleation activity by TMOD2 (31). Zoghbi and colleagues were the first to develop and characterize the *Tmod2* knockout mice and elegantly showed that its loss leads to novelty-induced hyperactivity, memory deficits in a fear conditioning task, and enhanced long term potentiation at hippocampal synapses (32). Congruently, recent human GWAS studies have shown that *Tmod2* haplotypes are associated with intellectual and cognitive impairments (33, 34). In human candidate gene studies for addiction traits, Gelernter and colleagues showed that *Tmod2* is associated with drug-dependent risk-taking behavior and proposed TMOD2-mediated remodeling of brain areas as a regulator of risky sexual behavior and drug intake (35). In several transcriptomics and proteomics studies, *Tmod2* mRNA and protein levels are differentially modulated following repeated methamphetamine (36, 37), alcohol (38), cocaine (39), or morphine (40) administration in animal models. These studies led a meta-analysis to suggest *Tmod2* is an addiction candidate gene (41). However, to date, there has not been a study that directly investigates a causal role of *Tmod2* in addiction phenotypes and provides mechanistic insights into its function in the reward circuit.

Based on recent human genetic data and our mining of the Knockout mouse project (KOMP) behavior phenotype data, we hypothesized that *Tmod2* is a regulator of addiction-related phenotypes. To test this, we carried out detailed addiction phenotype characterization for cocaine responses and find that sensitized response and intravenous self-administration (IVSA) acquisition are diminished in *Tmod2* knockout mice. Imaging and genomics data show reduced brain size but no major perturbances in gene co-expression networks, respectively. Finally, a detailed characterization of the intrinsic properties of cortical and striatal neurons at postnatal and adult stages show enhanced excitability. Synaptic events that are hallmarks of cocaine-induced synaptic plasticity in the nucleus accumbens medium spiny neurons are missing in the knockouts. Together, our data provide compelling evidence that the actin-regulating gene *Tmod2* plays a critical role in cocaine-induced behavioral adaptations that are critical for transition to addiction.

## Results

### Addiction-predictive behavior assessment using data in KOMP repository

A large body of literature suggests common underlying neurobiological pathways between addiction traits and other behavioral traits. Behavioral traits such as anxiety, novelty seeking, stress, and impulsivity, are predictive of alcohol- and drug-related phenotypes in humans and rodents. In humans, epidemiological studies show comorbidity between addiction phenotypes and sensation-seeking, impulsivity, and anxiety (42). These human studies are consistent with studies in experimental animals indicating that sensation seeking, novelty preference, and impulsivity predict psychostimulant self-administration (43).

Since KOMP uses metadata splits which can create imbalanced classes of controls and mutants, we reanalyzed the existing data using local controls (44). We analyzed the *Tmod2* phenotype data present in the public domain through the KOMP repository. Data from *Tmod2* knockout (n=16) and WT mice (C57BL/6NJ; n=68-92) that were tested concurrently ± 3 days were used for all the analyses. We specifically analyzed the results obtained from open field assay (OFA), light-dark box assay (LD), hole board (HB), paired-pulse inhibition test (PPI), and rotarod test (RR). All data are included as supplementary materials (Supplemental Table S1).

#### Hyperactivity and low anxiety in novel environment

Open-field behavior measures general locomotor activity, exploratory drive, anxiety-related behavior, and habituation learning. The mice were allowed to explore the open field arena, undisturbed, for 20 min and the activity was statistically compared in four, 5 min bins (Fig 1A1). *Tmod2* KO mice exhibit general hyperactivity that persisted for 20 minutes with a sign of habituation as seen by the bin-to-bin decline in distance traveled in 20 min. *Tmod2* KO mice show persistent hyperactivity compared to WT with increased number of breaks in all four, 5 min bins (Fig 1A1, genotype, F(1,328)=180, p<0.0001; time-bins, F(3,328)=22.43, p<0.0001; genotype X time-bins, F(3,328)=10.05, p<0.0001). Consequently, total time spent mobile was also significantly elevated in *Tmod2* KO (Fig 1A5, t=7.973, p<0.0001). *Tmod2* KO mice spend more time in the center (Fig 1A3, t=6.662, p<0.0001) and exhibit increased percent locomotor activity in the center of open field arena representing lower anxiety levels compared to WT (Fig 1A2, t=10.1, p<0.0001). Rearing, a risk assessment and investigative behavior (45), was reduced in KOs (Fig 1A4, t=2.055, p=0.0431). Therefore, the behavioral assessment of *Tmod2* KOs in an open field suggests a novelty-induced locomotor activity with decreased anxiety levels and diminished risk assessment or an increased propensity for risk-taking behavior.

**Figure 1.**
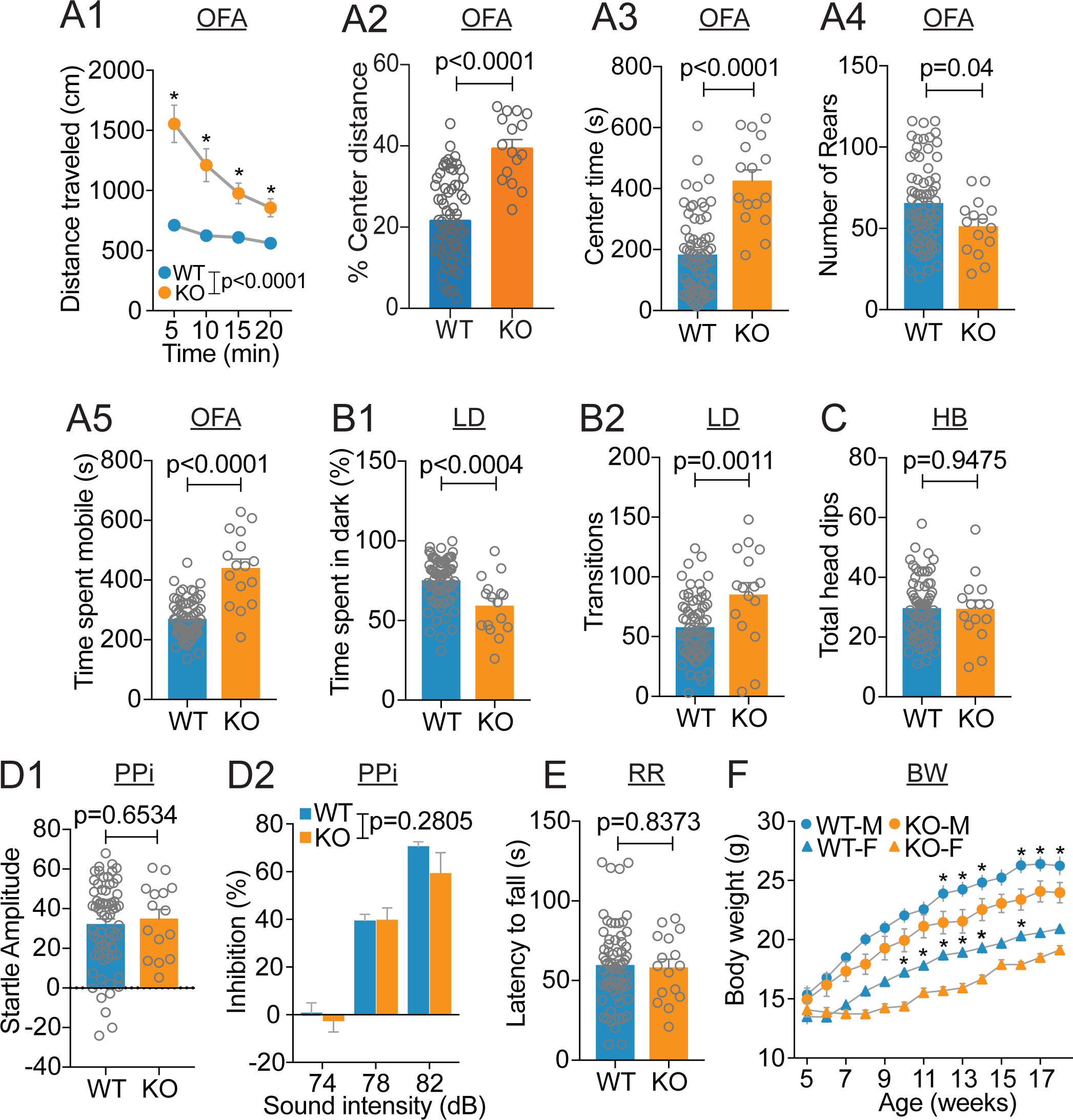
Behavioral characterization of *Tmod2* KO using publicly available data in KOMP repository shows specific deficits in hyperactivity and anxiety phenotypes. (A1-A5) Open-field assay results; (A1) Distance travelled in a 20 min exposure to standard open-field arena in four, 5 min. (*Bonferroni’s multiple comparison post hoc test detected significant differences (p<0.0001) between WT and KO mice.) (A2) percentage of distance travelled in center and (A2) mean total time spent in center. (A4) Number of rear postures attained. (A5) Total time spent mobile. (B1-B2) LD box results; (B1) time spent in dark side of the LD. (B2) Number of transitions between light and dark regions. (C) 20 min exposure to hole-board assay. (D1-D2) Prepulse inhibition assay results; (D1) startle amplitude and (D2) percentage inhibition of reaction to a subsequent stronger acoustic stimulus. (E) Rotarod performance test. (F) Only those mouse that has body weight data for all the 14 weeks were taken for analysis (WT, n=30, 15 males, 15 females; KO, n=16, 8 males, 8 females). Body weight growth curve (*significantly different from same-sex *Tmod2* KO, p<0.05). Error bar represents SEM. OFA, open field assay; LD, light-dark box; HB, hole-board assay; PPi, paired-pulse inhibition; RR, rota-rod test; BW, body weight growth curve.

#### Low anxiety and risk taking behavior towards aversive context

Behavioral assessment was carried out in a standard light-dark box with 2 zones with different light intensity. A brightly lit area is aversive and mice prefer the dark side. Exploration of the light side, as measured by the amount of time in the dark compared to the light side and the number of transitions between the two sides, is a form of risk-taking and anxiety behavior (46). *Tmod2* KO spent significantly less time in dark compared to WT (Fig 1B1, t=3.72, p=0.0004) and had a greater number of transitions between light and dark zone (Fig 1B2, t=3.382, p=0.0011). This assay provides complementary evidence of the novelty-induced hyperactivity, increased risk-taking, and low anxiety behavior in *Tmod2* KO that was seen in the open field assay.

#### Intact general exploratory behavior, sensory motor gating, and motor coordination and learning

Even though *Tmod2* KO appears to be hyperactive and less anxious in a novel environment, other parameters such as general exploration, sensory-motor gating, and motor learning are intact. General exploration is measured using the hole-board apparatus that consists of a floor with multiple holes (16 in total) and elicits an exploratory head-dipping exploratory behavior. Total head dips in a 20 min window were found to be comparable in *Tmod2* KO and WT (Fig 1C, t=0.06607, p=0.9475), indicating no deficits in general exploration behavior in *Tmod2* KO.

Sensory-motor gating is measured using paired-pulse inhibition protocol. This behavioral assay was carried out in a sound-attenuated chamber. Startle response amplitude (Fig 1D1, t=0.4508, p=0.6534) and percent inhibition to an increasing levels of pre-pulse sound prior to aversive sound stimuli were comparable in KOs and WT (Fig 1D2, genotype, F(1,240)=1.17; p=0.2805; sound intensity, F(2,240)=80.02, p<0.0001; genotype X sound intensity, F(2,240)=0.6452, p=0.5255), indicating unperturbed sensory motor gating in *Tmod2* KO. Effects of *Tmod2* deletion on motor-learning and coordination was assessed using the rotarod test. In this test, motor-learning is tested by calculating mean latency to fall in 5 consecutive trials on accelerating cylindrical rod. *Tmod2* KO mice show no deficits in motor learning tasks and their performance is comparable to their WT counterparts (Fig 1E, t=0.206, p=0.8373).

#### Decreased body weight which is slightly biased towards loss of lean mass

KOMP phenotyping pipeline also collects weekly measurements of body weight between 5^th^-18^th^ weeks of age. *Tmod2* KO males and females show a significant decrease in body weight relative to WT that is more pronounced in late adulthood (10-12 weeks of age) (Fig 1F, genotype, F(1,42)=11.59, p=0.001; age X genotype, F(13,546)=10.38, p<0.0001; age X sex, F(13,546)=19.67, p<0.0001; genotype X sex, F(1,42)=0.001, p=0.9741). Further exploration of body composition data from microCT scans revealed an increased lean body mass (Supplementary Fig S1E, t=2.437, p=0.017), suggestive of decreased fat mass (Fig S1F, t=1.719, p=0.0893) and a significant reduction in body length (Fig S1D, t=4.43, p<0.0001) in *Tmod2* KO. Therefore, an age-dependent decrease in body weight in *Tmod2* KO is due to a modified lean vs fat mass ratio and hindered overall body growth or a combination of both. These data indicate that the *Tmod2* KO mice are approximately 22-28% smaller starting at 10-11 weeks of age. Although smaller, these mice had no overt phenotype(s) and did not show any signs of illness or poor health. Combined, KOMP data demonstrate specific behavior signatures in *Tmod2* KO that have been shown previously to be predictive of addiction (43, 47), while several other parameters were found unchanged when compared to WT counterparts (see also Supplementary Fig S1).

### Comprehensive phenotyping for addiction-relevant behavior

Based on KOMP results we generated a *Tmod2* KO colony and carried out addiction phenotyping. Prior to any behavioral analysis, we confirmed that TMOD2 was absent through western blot analysis. The results confirmed the absence of TMOD2 protein in the mutant’s cortical punches (Fig 2A). In addition, later genomics analysis showed reduced levels of *Tmod2* transcript (Fig 3D).

**Figure 2.**
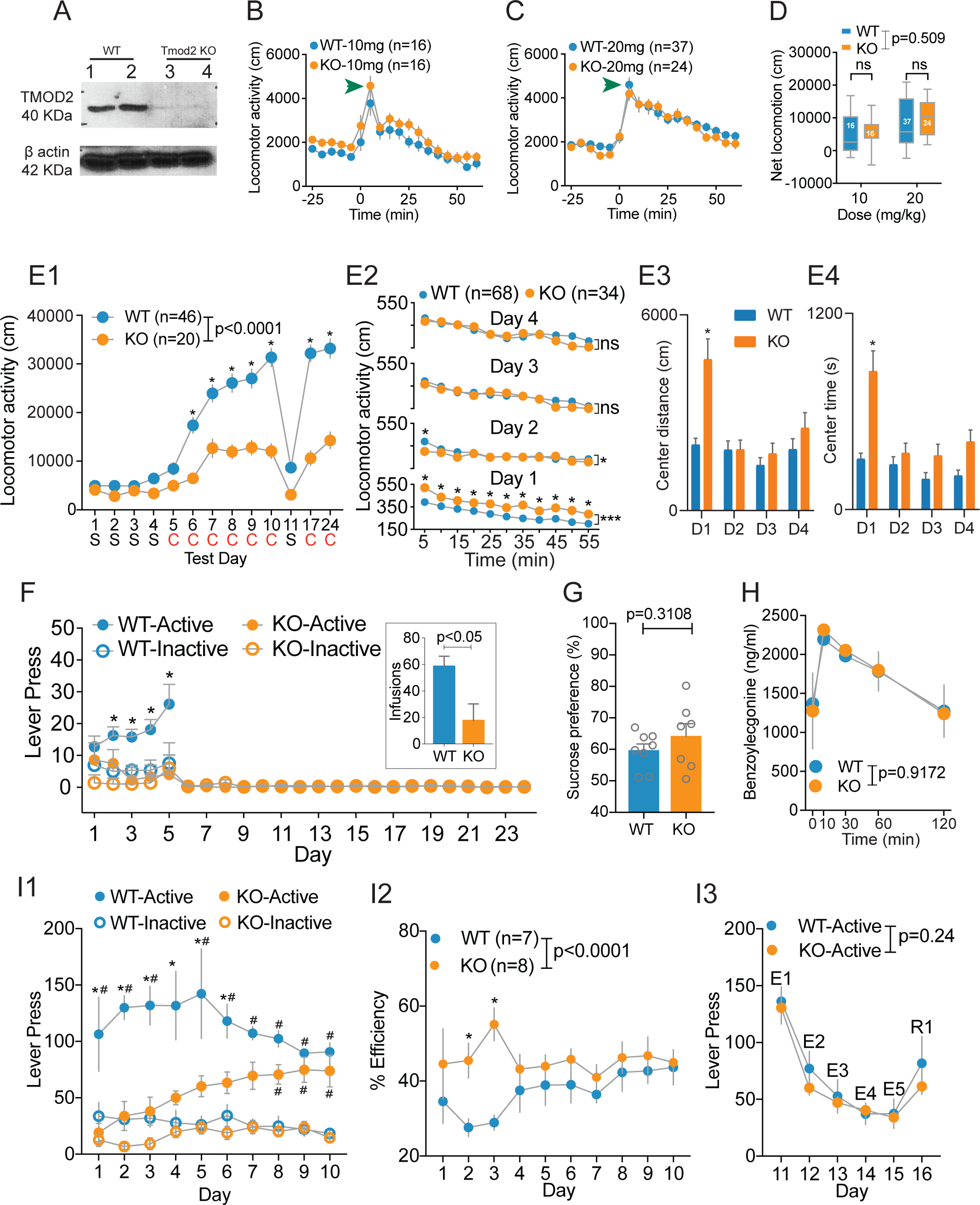
Comprehensive addiction phenotyping in *Tmod2* KO reveals deficits in cocaine sensitization and IVSA. (A) Qualitative assessment of absence of TMOD2 protein in cortical lysate from KO brain. (B) Locomotor response following acute injections of low (10 mg/kg i.p.) and (C) high (20 mg/kg i.p.) doses of cocaine. Green arrow head indicates locomotor artifact due to injection procedure. (D) Net locomotor activity following low and high dose of cocaine. (E1-E4) Results obtained from locomotor sensitization protocol. (E1) Graph’s x-axis show day and corresponding treatment (S, saline; C, cocaine) on that day. (E2) Day 1-4 baseline locomotor activity. (E3) Mean center distance travelled and (E4) time spent in center during baseline period of Day1-4. (F) Active and inactive lever presses on day 1-24 of cocaine IVSA testing. Inset show number of cocaine infusions. (G) Sucrose preference. (H) Cocaine metabolite benzoylecgonine concentration in plasma at different time points following cocaine (20 mg/kg) injections. (I1-I3) Operant conditioning using palatable food as a reinforcement resulted in (I1) Active and inactive lever presses; (I2) Total sucrose pellets obtained by total number of active and inactive lever presses is calculated as percent efficiency. (I3) Extinction sessions (E1-E5 on day 11-15) and reinstatement (R1 on day 16) of reward behavior. Error bar represents SEM. ns, not significant. *Bonferroni’s multiple comparison post hoc test detected significant differences (p<0.01) between WT and KO mice. ***main effect of genotype detected using 2way ANOVA (F10, 1100) = 25.10, p<0.0001). ^#^Significant difference between inactive and active lever press within same genotype (p<0.01).

**Figure 3.**
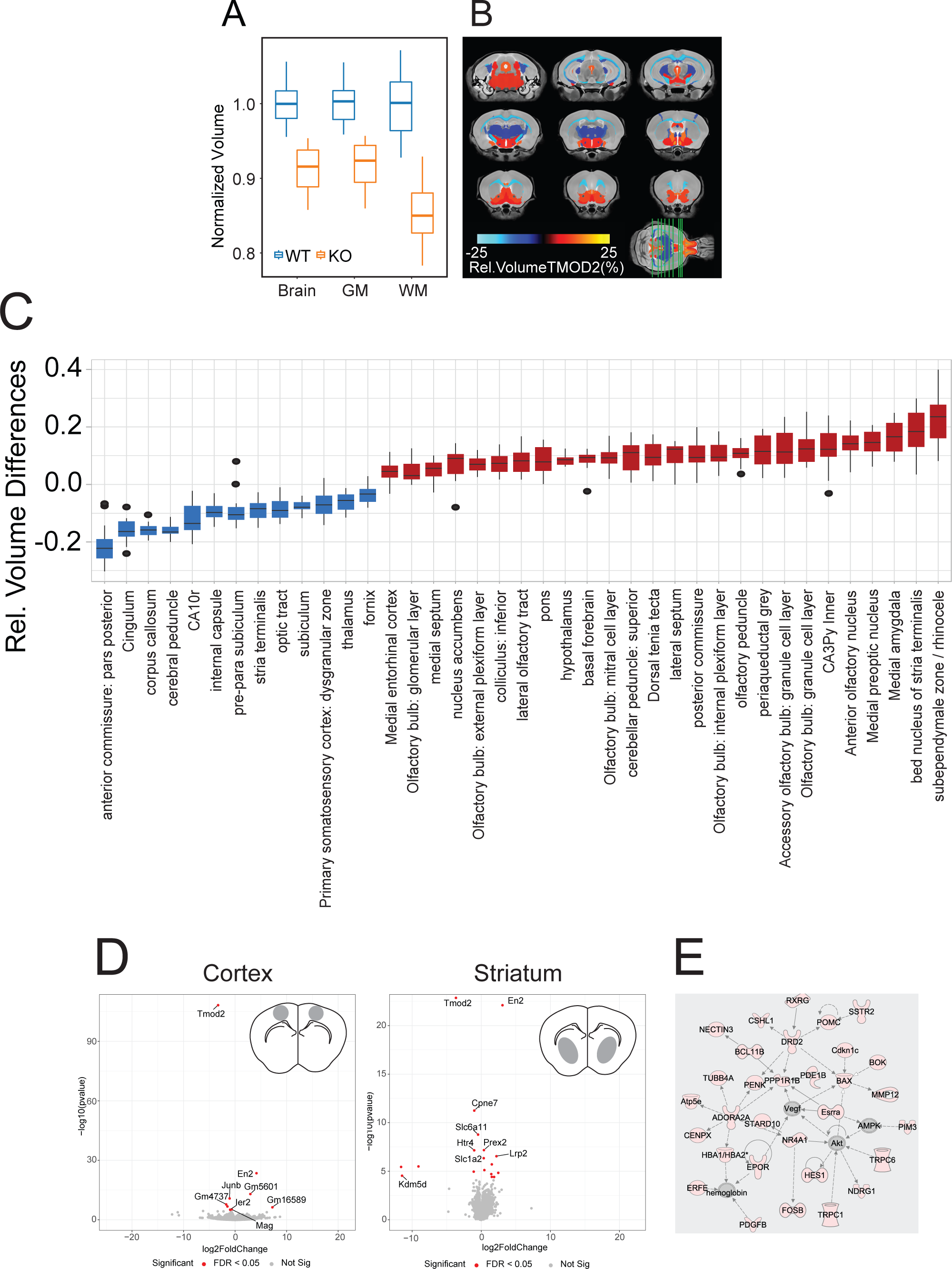
MRI and bulk RNAseq analysis on WT and KO brains. (A) Normalized brain volume comparison. (B) KO strain differences based on relative volumes. Color overlays indicate volume changes referenced to the WT. (C) Relative volume differences of KO brain regions. Blue and red boxes indicate relative decrease and increase in volume of labelled brain region. Most of the blue boxes are associated with white matter regions in the brain. (D) Schematic of cortical punches obtained from M1 sub-region and dorso-ventral striatum (grey highlighted region) along with volcano plots of cortical and striatal differentially expressed genes in KO relative to WT. (E) IPA analysis of *Tmod2* KO primary and secondary motor cortex transcriptome. Most component of this gene regulatory network are reduced in expression as indicated by peach-colored modules.

#### Unaffected acute response to low and high doses of cocaine

Separate groups of WT and KO mice were acutely injected with cocaine (10 or 20 mg/kg) and their locomotor response was analyzed 30 min before and 1h following injections. (Fig 2B-C). For statistical comparison, the net locomotor response was calculated by subtracting the total locomotor activity in a 30 min period following cocaine injections with the locomotor activity during 30 min baseline. Net locomotor response was comparable between mutants and WT for both doses of cocaine indicating that the neural circuits that evoke acute drug-induced locomotor response is intact in *Tmod2* KO (Fig 2D, genotype, F(1,89)=0.509, p=0.4789; dose, F(1,89)=10.07, p=0.0021; genotype X dose, F(1,89)=0.08954, p=0.7655).

#### Cocaine-induced behavioral sensitization is impaired in Tmod2 KO

Repeated administration of psychostimulants such as cocaine evokes behavioral sensitization - a progressive and persistent amplification of locomotor response to the same dose of the drug. Sensitization is a hallmark of many drugs of abuse and is due to neuroadaptations in the reward circuit (48, 49). Although it is only partially predictive of self-administration, a sensitization paradigm is often used to gauge plasticity in the mesolimbic circuit (50). Our sensitization protocol consists of once-daily injections of saline for 4 consecutive days (test day 1-4), followed by 6, non-contingent, cocaine injections (15mg/kg, test day 5-10) and a saline injection on test day 11 (Fig 2E1). Since sensitization has been shown to be long-lasting (51), we carry out a cocaine challenge on test day 17 and 24, a period of 1 or 2 weeks later. Repeated administration of cocaine resulted in progressive amplification of locomotor responses in WT mice (Fig 2E1). Intriguingly, *Tmod2* KO had a significant attenuation of the cocaine-induced sensitization response compared to WT (Fig 2E1, genotype, F(1,806)=252, p<0.0001; drug, F(12,806)=51.42, p<0.0001; genotype X drug, F(12,806)=11.34, p<0.0001). *Tmod2* KOs show slight sensitized responses to the first three cocaine injections (day 5-7) with an increase in locomotor activity, but from the 4^th^ day onwards (day 8-10) and on cocaine challenge days (day 14 and 24), no further escalation in locomotor activity was observed (Fig 2E1). It is worth noting that a weak, drug-induced sensitization response is exhibited by *Tmod2* KO on day 5-7 with an early-achieved ceiling effect that stabilizes locomotor response from the 8^th^ day onwards and persists for rest of the protocol. Post hoc analysis confirmed the absence of any locomotor differences between mutant and WT to acute cocaine administration (Fig 2E1, day 5, 1^st^ cocaine administration, p=0.8109). We visually inspected the recorded videos for stereotyped behaviors, another form of behavioral sensitization (52, 53), but didn’t observed any cocaine-induced short or long-term effects in *Tmod2* KO mice. These data indicate an impaired behavioral sensitization response in *Tmod2* KO suggestive of compromised drug-induced neuroadaptations.

Multiple exposures to a similar environmental context attenuates novelty-induced locomotor activity and leads to habituation: a form of non-associative learning (54). In our analysis of sensitization data, we observed that *Tmod2* KO mice were not hyperactive in the open field upon repeated testing, indicating high rates of habituation. Pre-injection baseline periods (−60 to 0 min) of the first 4 saline days of behavioral sensitization paradigm were used to reconfirm the open-field phenotype observed in the KOMP analysis. *Tmod2* KO exhibited persistent, elevated locomotor activity” compared to WT on day 1 (Fig 2E2-Day 1, genotype, F(1,1100)=147.5, p<0.0001; time-bins, F(10,1100)=25.1, p<0.0001; genotype X time-bins, F(10,1100)=0.5143, p=0.8809). A main effect of genotype was also detected on day 2 (Fig 2E2-Day 2, genotype, F(1,1100)=4.995, p=0.0256; time-bins, F(10,1100)=6.166, p<0.0001; genotype X time-bins, F(10,1100)=1.007, p=0.4356) however, the post hoc analysis revealed that the time-bins significance was restricted to the first, 5-minute bin with WT having more locomotor activity compared to *Tmod2* KO (Fig 2E2-Day 2, Sidak’s multiple comparison, p=0.0245). On day 3 (Fig 2E2-Day 3, genotype, F(1,1100)=2.298, p=0.1298; time-bins, F(10,1100)=18.07, p<0.0001; genotype X time-bins, F(10,1100)=0.9725, p=0.4656) and day 4 (Fig 2E2-Day 4, genotype, F(1,1100)=0.5777, p=0.4474; time-bins, F(10,1100)=14.34, p<0.0001; genotype X time-bins, F(10,1100)=1.211, p=0.2794), a main effect of genotype and post-hoc time-bins comparison failed to detect any statistically significant difference between *Tmod2* KO and WT. Similarly, on day 1, center distance traveled (Fig 2E3, D1, t=5.386, p<0.001) and time-spent in the center (Fig 2E4, D1, t=6.559, p<0.001) are significantly higher in *Tmod2* KO recapitulating the results obtained from KOMP data. However, from the 2^nd^ day onwards (D2-D4, Fig 2E3-4), these anxiety parameters are comparable in both the genotypes, indicating an amplified habituation response to a less novel, familiar environmental context in *Tmod2* KO. This analysis demonstrates that the hyperactivity in *Tmod2* KO is a specific response towards environmental novelty that is only seen on the first day of the open field test and is not due to general hyperactivity. Thus, habituation, a form of non-associative learning is preserved and perhaps increased, in *Tmod2* KO. Combined, these data suggest that experience and pathway-specific neuroadaptations that mediate habituation are functional in *Tmod2* KO, while specific deficits are seen in neuroadaptations that mediate behavioral sensitization.

#### Failure to acquire cocaine self-administration

Although locomotor sensitization is an established model of drug-induced plasticity and neuroadaptations, it is only partially predictive of voluntary drug intake (50). Goal-directed behavior and motivation to procure drug is conventionally tested by intravenous self-administration assay (IVSA), the gold standard technique in the addiction field (55). Therefore, we assessed cocaine acquisition and dose-response in *Tmod2* KO and WT using the IVSA paradigm.

To assess cocaine IVSA performance, we performed repeated measures or one-way ANOVA using the number of lever presses or number of infusions as dependent measures. Genotype (*Tmod2* KO, WT) and sex were between-subject factors. Session (day 1 - 5) was a within-subjects factor. When lever presses were used as the dependent measure, lever (active, inactive) was used as a second within-subjects factor. WT mice rapidly learned to lever press for cocaine (Figure 2F) as indicated by a significant increase in active lever presses across sessions (day 1 vs day 5: p = .05) and a significant dissociation between the active and inactive levers (active vs inactive on s5: p = .002). In contrast, *Tmod2* KO mice failed to acquire cocaine infusions as indicated by a failure to significantly increase active lever presses across sessions (day 1 vs day 5: p = 0.67) and a failure to dissociate between the active and inactive lever (active vs inactive on day 5: p = 0.82). This lever pressing pattern resulted in WT mice infusing significantly more cocaine across the first five sessions relative to *Tmod2* KO mice (Fig 2F inset, p<0.05). Lever pressing was mediated by strain and lever as indicated by significant main effects of strain [F (1, 26) = 5.00, p = .03] and lever [F (1, 26) = 12.84, p = .001], as well as a significant strain X lever interaction [F (1, 26) = 4.64, p = .04]. Interestingly, active lever press were comparable on day 1 indicating that general exploration and responses to operant cues are able to evoke behavior directed to the levers in *Tmod2* KO. However, either neutral or aversive outcome of active lever press rapidly extinguishes the future responses towards the active lever.

Comparison of WT and *Tmod2* KO mice for cocaine IVSA was limited to the acquisition phase because WT mice stabilized at the 1.0 mg/kg/infusion acquisition dose and advanced to the dose-response curve stage by session five. *Tmod2* KO mice were tested for a total of 24 sessions (Fig 2F) to test if they could learn to acquire cocaine if sufficient exposure was provided. During these 24 sessions, *Tmod2* KO mice failed to significantly increase response on the active lever and failed to dissociate between the active and inactive lever as indicated by nonsignificant main effects of lever (p = 0.95) and session (p = 0.24) as well as a nonsignificant lever x session interaction (p = 0.33). Since the mice failed to acquire, cocaine IVSA testing was terminated for *Tmod2* KO mice after session 24. IVSA results demonstrate that the dose which is reinforcing to WT mice is not reinforcing to KO mice and indicates either general reward deficits, learning deficits in an operant task, or both. Additionally, modified cocaine metabolism could also contribute to lack of acquisition and low locomotor sensitization in *Tmod2* KO.

#### Intact motivation towards natural reward

In order to investigate the responsivity to natural reward and to rule out a general deficit in reward-based learning, we subjected mice to a two-bottle sucrose preference test. Both strains show comparable motivation and preference towards 4% sucrose solution provided in the sucrose preference test (Fig 2G, t=1.055, p=0.3108). Therefore, inability of *Tmod2* KO to self-administer cocaine is not because of a general deficit in reward-sensing and responding neural circuitry.

#### Cocaine metabolism is unaffected in Tmod2 KO mice

*In vivo* cocaine metabolism is an important determinant of its pharmacokinetics and tissue bioavailability that could modulate a drug-induced addiction-like phenotype. To test cocaine metabolism in these mice we carried out a biochemical analysis for primary cocaine metabolite benzoylecgonine (56) by injecting mice with cocaine (20mg/kg) and collecting blood from the submandibular vein at 0, 10, 30, 60, and 120 min following injections. No significant differences in the level of the cocaine metabolite, benzoylecgonine, was observed between *Tmod2* KO and WT mice for any of the sampled time points, indicating that cocaine metabolism is not affected in these mutants (Fig 2H).

#### Delayed learning ability in operant assay

Since *Tmod2* has previously been shown to regulate learning and memory deficits in mice (32) and has been linked to intellectual deficits in human GWAS studies (33, 34), we tested whether the KO mice were capable of learning an operant task. Operant conditioning relies on associative learning and therefore the impairment to acquire cocaine in the IVSA paradigm could be due to learning deficits. We investigated operant learning in *Tmod2* KO mice for palatable food reward as a reinforcement stimulus. *Tmod2* KO mice took significantly more number of trials to learn the association between active lever and palatable food reward, and to discriminate between the active and inactive lever (Fig 2I1). In contrast, WT mice learned to discriminate between the active and inactive lever by the end of day 1 and demonstrates stronger associative learning than *Tmod2* KOs (Fig 2I1, genotype & lever, F(3,242)=117.9, p<0.0001; day, F(9,242)=0.7612, p=0.6525; genotype & lever X day, F(27,242)=1.347, p=0.1244). In *Tmod2* KO, statistically significant discrimination between the active and inactive lever emerges on day 8, and from day 7 onwards, active lever presses by KO and WT are comparable (Fig 2I1). Surprisingly, operant efficiency (number of sucrose pellets obtained divided by total number of active and inactive lever press) was slightly better in *Tmod2* KO. We hypothesize that this is due to compulsive lever pressing by WT mice during initial trials (Fig 4F1, day 1-5) which ignore the 20 s timeout period between two successful (paired with reward) active lever presses (Fig 2I2, genotype, F(1,115)=19.37, p<0.0001; day, F(9,115)=0.7663, p=0.6477; genotype X day, F(9,115)=1.554, p=0.1375). On extinction sessions (Fig 2I3, E1-E5 on day 11-15) during which neither reward nor operant cue (cue light off) were presented, both *Tmod2* KO and WT exhibited a comparable, rapid decline in the number of lever presses that was previously paired with reward indicating that extinction-induced associative learning is intact in both the genotypes (Fig 2I3, genotype F(1,66)=1.405, p=0.2401; day, F(5,66)=18.72, p<0.0001; genotype X day, F(5,66)=0.2839, p=0.9203). Rebound of behavior directed towards active lever with one single reinstatement session was also comparable between *Tmod2* KO and WT mice (Fig 2I3, R1, day 16, Sidak’s post hoc test, p=0.79). Thus, as shown previously, *Tmod2* KO mice have a deficit in learning (32). However given enough time and trials, *Tmod2* KO mice learn an operant tasks. In trials with food reward, *Tmod2* KO mice were indistinguishable from WT mice from day 7, whereas with cocaine as a reinforcement *Tmod2* KO failed to learn even after 24 day trial, which leads us to conclude that there is a deficit in cocaine-induced reward learning in *Tmod2* KO mice.

#### Reduced brain volume and alteration of gene expression networks in Tmod2 Knockout mice

Since neuron-specific TMOD2 regulates actin stabilization and plays an important role in dendritic arborization, and synapse formation (27, 28), we investigated if regional and gross morphological changes are seen in *Tmod2* KO brains. We carried out MRI analysis, a well-established diagnostic and translational tool in clinical and preclinical research, and discovered that whole-brain volume is significantly decreased in *Tmod2* KO relative to the WT mice (Fig 3A-B, −8.8%, p=1.5×10^-9^). The extent of volume loss appears to be greater in white matter, as the overall changes for gray and white matter were −8.2% (p=4×10^-9^) and - 14.7% (p=7×10^-12^) respectively (Fig 3A). The volumes of 182 structures (using combined left-right volumes) were computed for all animals. While the 20-25% decrease in body weight in 10-11 week-old *Tmod2* KO (Fig 1F) may explain some of this change, proportional weight change is not generally a predictor of proportional brain volume change in adult rodents (57, 58).

For the absolute volume analysis, a total of 109 structures were observed to be significantly different for the *Tmod2* KO mice (q<0.05), all but one of which were volume decreases. Prominent changes included regions of the cerebellum, much of the cortex, the corpus callosum, the thalamus, the hippocampus, and, with relevance to the mesolimbic system, the striatum and midbrain (summarized in Supplemental Table S2). However, given the overall difference in brain size, we also normalized results by expressing structure volume as a percentage of whole-brain volume, and then comparing results. A total of 39 brain structures showed a significant difference in relative volume (Fig 3B). Several of the regions decreased in absolute size were still observed to be decreased in this analysis, especially the white matter regions (Fig 3B, 3C blue bars), but several structures also appeared relatively larger (Fig 3B, 3C red bars), including the hypothalamus, olfactory bulbs, nucleus accumbens and basal forebrain.

We next, attempted to detect any molecular differences that may have resulted from the deletion of *Tmod2* gene. Specifically, we hypothesized differences in cell type constituency, cell loss, neuroinflammation, neurodegeneration, or general developmental deficits that may be cell-type specific and thus, would lead to changes in gene co-expression networks that could be detected by monitoring of genome-wide gene expression (59, 60). We carried out RNAseq analysis from striatum and cortex of naïve adult mice (3M/3F) for *Tmod2* KO and WT animals. A PolyA enriched RNA sequencing library was generated at a depth of 40-50 million reads. To determine which gene regulatory networks are perturbed in *Tmod2* mice, we carried out RNA-seq analysis from motor cortex and striatum of naïve 10 week old mice. Gene sets found to be misregulated (Figure 3D) were further analyzed by Qiagen Ingenuity Pathway Analysis software (62). We observed numerous disease categories were significantly enriched in both tissues, including neurological disease, behavior, and organismal injury (177 genes, p = 1.40E^-02^ - 4.46E^-19^) and neurological disease and nervous system development (124 genes, p = 1.4 x 10^-2^ to 4.6×10^-17^) (Figure 3). The top molecular function perturbed was cell morphology (135 genes, p 1.30×10^-02^ - 1.07×10^-13^) and top physiological function of these genes was nervous system development (177 genes, p = 1.40×10^-02^ - 4.46×10^-19^). Prominent perturbations were observed in the expression of genes that regulate cell structure and neurotransmission (e.g., DRD2, POMC, serotonin receptors) (Fig 3E). This gene expression analysis points towards a developmental defect in both the cortex and striatum of Tmod2 *mice*, consistent with the massive volume decreases seen throughout the brain in MRI scans of *Tmod2* animals, something previously linked to addiction (63). As a control analysis, we compared striatum and cortex gene expression in WT animals and found over 7439 genes (Supplemental Table S3) that were differentially expressed in these regions (Fig 3E, FDR<0.05). These genomics results indicate that there are specific changes in gene coexpression network that support the developmental and structural deficits observed in *Tmod2* KO. Thus, structural and transcriptomic changes in Tmod2 KO suggest functional differences as an underlying cause of emotionality and addiction-relevant phenotype in these mutants. This led us to hypothesize that the observed behavioral changes are due to functional differences that may be detectable using electrophysiological methods.

### Enhanced cortical excitability during development and adulthood

Since TMOD2 regulates dendritic arborization (27, 64), synapse density and maturation, and synaptic plasticity (11, 27, 28), we hypothesized that the firing properties of neurons might be modified due to altered synaptic connectivity. To test this, we harvested cortical cells from P0-P1 pups and quantified the spontaneous firing features of cultured neurons on MEA plates. We also tested the effect of GABA-A receptor and AMPA receptor antagonism on spontaneous activity (Fig 4B). MEA captures *in vitro* real-time electrophysiological activity and interconnectivity between cultured neurons permitting recreation of complex firing patterns and detection of deficits in a high-throughput fashion. *Tmod2* protein expression was detected in the rat brain as early as embryonic day 14 (E14) and attained adult-level expression by E19 (30). Embryo imaging data collected on *Tmod2* KO by KOMP phenotyping reveals robust, nervous system-specific LacZ expression driven by the *Tmod2* promoter at E12.5 (Fig 4A). This early developmental expression of *Tmod2* in mouse embryos suggests a critical function that could be compromised in *Tmod2* KO. *Tmod2* KO cortical neurons discharged at a higher frequency in a drug-free condition (Fig 4C-E1, t=3.547, p=0.0045). Picrotoxin, a potent GABA-A receptor blocker, failed to affect the firing rate in *Tmod2* KO cultures, whereas WT neurons respond with a three-fold change in firing rate (Fig 4E2, t=2.899, p=0.0327). This could be due to fewer functional, membrane-bound GABA-A receptors in *Tmod2* KO neurons or the absence of presynaptic GABAergic innervations or both. Absence or decrease in inhibitory tone to cultured neurons could also explain increased firing rate in *Tmod2* KO cortical cultures. Similarly, *Tmod2* KO neurons produce significantly more bursts in the drug-free baseline condition (Fig 4F1, t=4.392, p=0.0011). Picrotoxin application resulted in an eight-fold increase in burst number in WT neurons and has no effect on *Tmod2* KO neurons suggesting altered GABA receptor availability or GABAergic synapses or both (Fig 4F2, t=4.538, 0.0094). NBQX application prevents bursting in WT cortical neurons while the *Tmod2* KO neurons showed a similar tendency, but did not reach our statistical cutoff (Fig 4F2, t=2.391, p=0.0863). Synchronicity among cultured neurons to fire simultaneously is significantly elevated in *Tmod2* KO neurons (Fig 4G1, t=5.133, p=0.0003). PTX application resulted in a four-fold increase in firing synchronization of WT cortical neurons and had no effect on *Tmod2* KO neurons (Fig 4G2, t=4.947, p=0.003). As anticipated, NBQX application mitigates the synchrony index in WT neurons but not in *Tmod2* KO (Fig 4G2, t=3.16, p=0.0134). These results clearly demonstrate that *Tmod2* KO neurons in culture exhibit enhanced cortical excitability along with modified excitation/inhibition ratio during early developmental stages.

**Figure 4.**
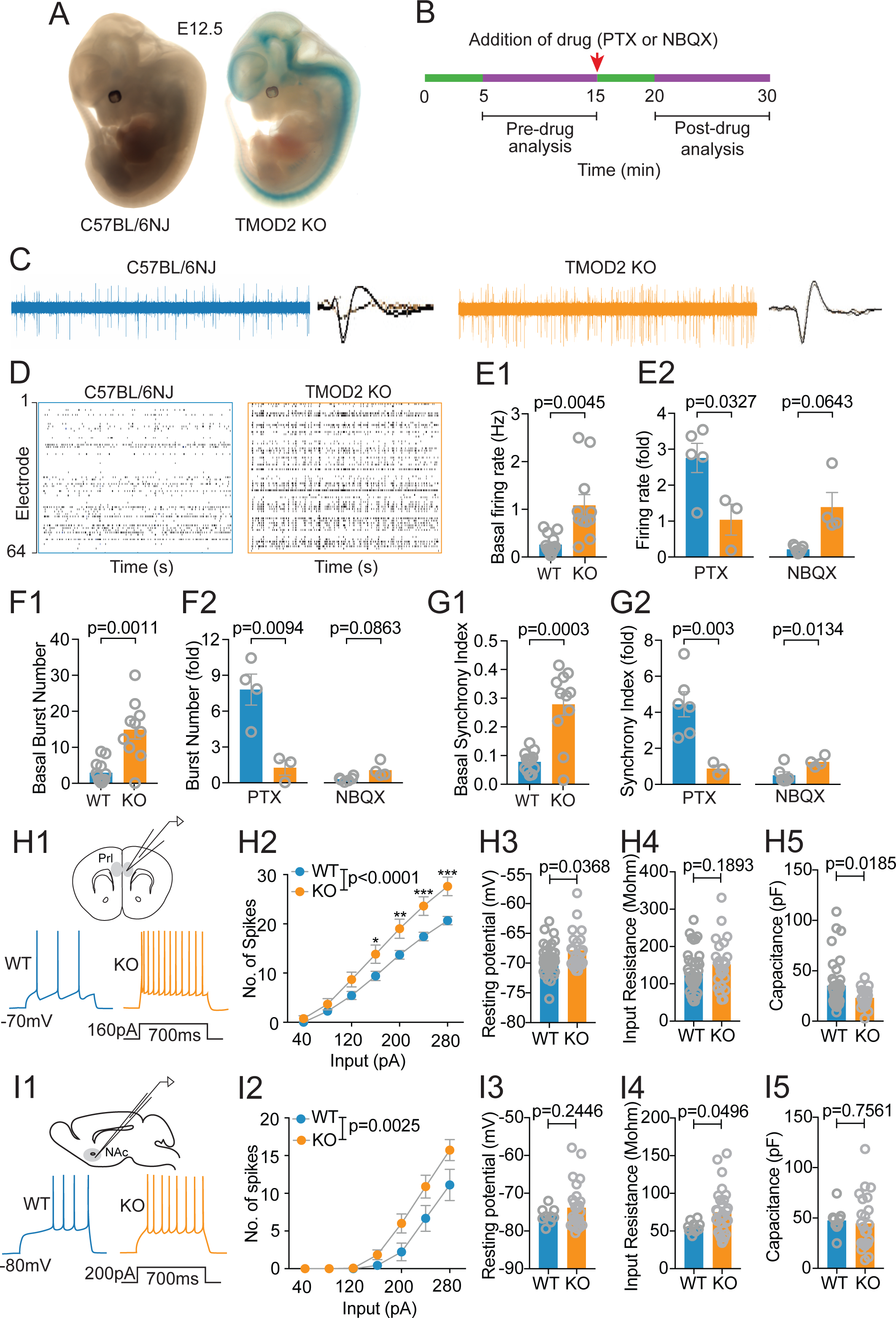
Characterization of spontaneous firing features of early postnatal cortical neurons and intrinsic properties of reward regions in adult brain. (A) LacZ expression in KO embryo (E12.5) driven by *Tmod2* promotor and absence of expression in WT. (B) Timeline of multielectrode array (MEA) recordings of cultured cortical neurons. (C) Raw traces of spiking activity and offline spike sorting captured by MEA electrodes. (D) Representative raster plots of spontaneous spiking activity in WT and KO neurons across 64 electrodes. (E1) Baseline spiking activity obtained from pre-drug window. (E2) Spiking activity following PTX and NBQX application in bath. (F1) Number of burst detected in 10 min pre-drug period and (F2) following PTX and NBQX application. (G1) Synchrony index; (G2) synchrony index following PTX and NBQX application. (H1-H5) Intrinsic properties of adult, prelimbic pyramidal neurons. N= 23-38 neurons per genotype; 3-4 mice per genotype (H1) Schematic of whole-cell recordings performed in 5^th^ layer of prelimbic cortex along with representative action potentials recorded by applying 160 pA for 700 ms. (H2) Mean number of spikes for a given (x-axis) magnitude of injected current. (H4) Input resistance. (H5) Whole cell capacitance. (I1-I5) Intrinsic properties of adult, accumbens shell medium spiny neurons (N = 9-28 neurons per genotype; 3-4 mice per genotype). (I1) Schematic of recorded brain region and samples traces at 200 pA. (I2) Mean number of spikes for a given (x-axis) magnitude of injected current. (I3) Resting membrane potential. (I4) Input resistance. (I5) Whole cell capacitance. Error bar represents SEM. *p<0.05, **p<0.01, ***p<0.001. E12.5, embryonic day 12.5. PTX, GABA-A receptor blocker. NBQX, AMPA receptor antagonist. Prl, prelimbic. NAc, nucleus accumbens.

To test whether neuronal excitability seen in early postnatal neurons persists in adult *Tmod2* KO, we compared intrinsic properties of prelimbic cortical (Fig 4H1-5) and accumbens shell neurons (Fig 4I1-5) in KO and WT using whole-cell recording in acute brain slices. Naïve mice were sacrificed and brain slices containing prelimbic and accumbens region neurons were recorded. *Tmod2* KO cortical neurons fired more action potential spikes in response to depolarizing current steps than WT neurons (Fig 4H1-2). *Tmod2* KO neurons showed slightly higher resting membrane potentials than WT neurons (Fig 4H3), but no change in input resistance or whole cell capacitance (Fig. 4H4-H5). The action potential threshold (39.51 ± 0.46 mV in WT and -38.6 ± 0.53 in *Tmod2* KO; Supplementary figure S3A1) and after-hyperpolarization amplitude (12.11 ± 0.62 mV in WT and 13.48 ± 0.76 mV in *Tmod2* KO; Supplementary figure S3A2) are statistically comparable between genotypes indicating functional similarity of voltage-gated sodium channels and potassium channels. *Tmod2* KO accumbens shell neurons are also more excitable than WT (Fig 4I1-2), but no change in resting membrane potential was observed (Fig 4I3). The input resistance is slightly higher in *Tmod2* KO neurons (Fig 4I4), and whole cell capacitance is comparable between both genotypes (Fig 4I5). The action potential threshold (−35.69 ± 0.98 mV in WT, −35.67 ± 0.59 mV in *Tmod2* KO; Supplementary figure S3B1) and after-hyperpolarization amplitude (12.05 ± 1.03 mV in WT and 12.07 ± 0.52 mV in *Tmod2* KO; Supplementary figure S3B2) are statistically comparable between genotypes. Therefore, *Tmod2* deletion results in an increase in excitability of prelimbic pyramidal neurons, and medium spiny neurons of accumbens shell, indicative of its role in determining the firing features of neurons and their interconnectivity.

### Distinct naïve and cocaine-induced synaptic properties

Action-potential independent, spontaneous release of neurotransmitters from presynaptic neurons and measurement of the miniature excitatory or inhibitory postsynaptic currents is a standard method to assess the baseline and drug-induced alteration in synaptic coupling between neurons. Non-contingent cocaine administration induces widespread modifications of synaptic properties in reward-related brain regions, triggering maladaptive reward learning (1). These synaptic alterations have been extensively characterized in nucleus accumbens and establish both effect and causality of cocaine abuse (1). *Tmod2* KO mice have an attenuated locomotor sensitization response to repeated cocaine administration and perform poorly in an IVSA paradigm. We hypothesized that drug-induced synaptic changes are either missing or weak in *Tmod2* KO resulting in a drug-resistant phenotype. Mice were either treated with a once-daily injection of saline or cocaine for 5 consecutive days, sacrificed following 2-5 day of abstinence and brain slices containing accumbens region neurons were recorded (Fig 5B). We compared mEPSC (Fig 5C1) and mIPSC (Fig 5D1) in medium spiny projection neurons of accumbens shell (Fig 5A) in drug-naïve (saline injected) and drug-experienced (cocaine injected) mice using *ex vivo* whole-cell recording in brain slices. Mean amplitude of mEPSCs is significantly elevated in naïve *Tmod2* KO compared with naïve WT neurons (Fig 5C2), which suggests an increased number or function of AMPA receptors on patched accumbens neurons (65). Cocaine treatment increases mEPSC frequency in WT but not *Tmod2* KO neurons (Fig 5B3), and suggests that repeated cocaine exposure fails to induce synaptic changes typically associated with chronic cocaine exposure (66). No significant effect of treatment or genotype was observed in either mEPSC rise or decay time which indicates a lack of alterations in the postsynaptic properties of AMPA receptors (Fig 5C4-C5).

**Figure 5.**
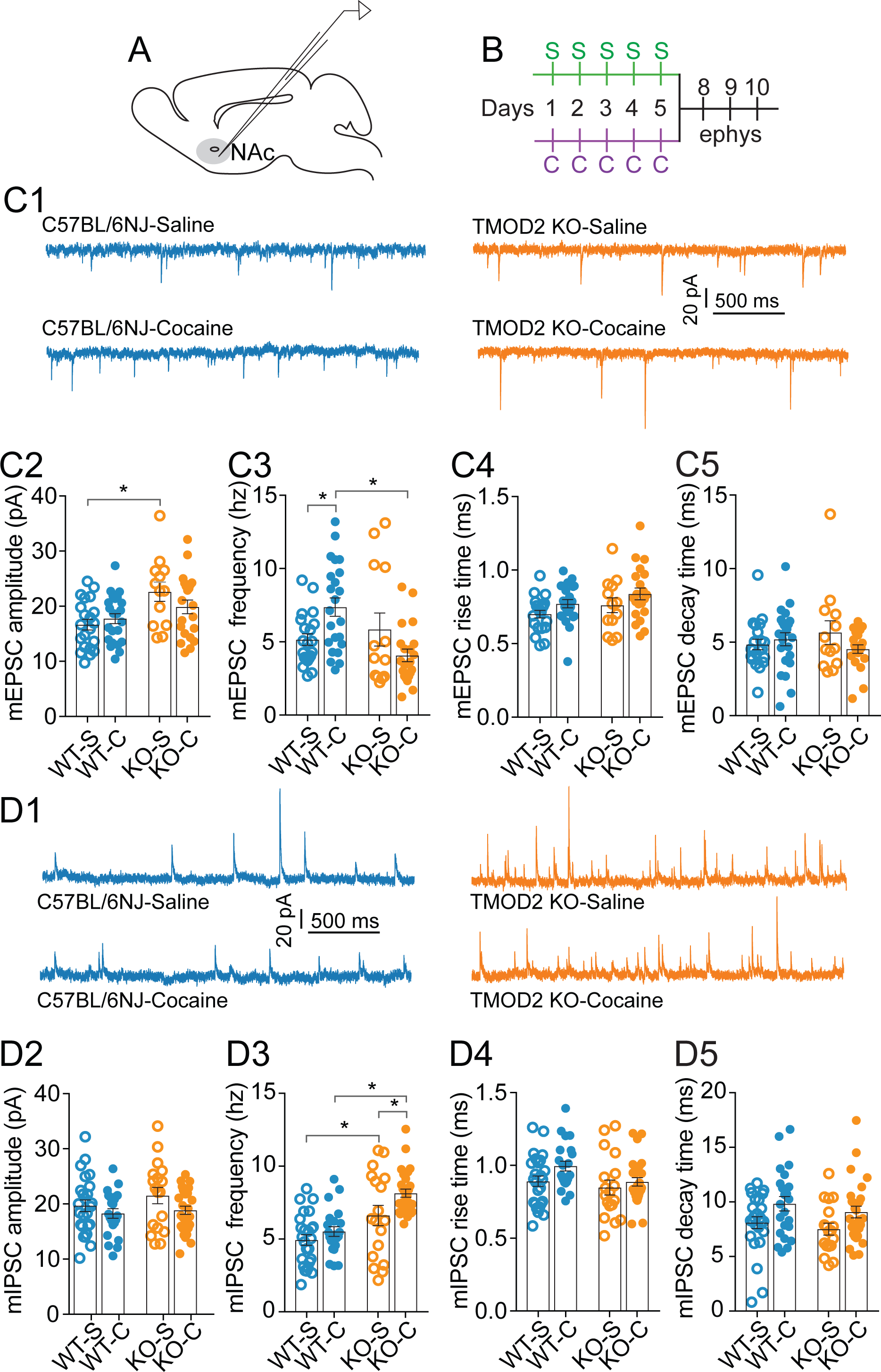
Electrophysiological characterization of glutamatergic and GABAergic synaptic transmission in accumbens shell medium spiny neurons. (A) Schematic of sagittal section of mouse brain containing recorded nucleus accumbens shell subregion. (B) Timeline of sensitization protocol in two groups. (C1) representative traces showing mEPSCs at holding potential of −80 mV in the presence of 0.5 µM TTX in saline (top) or cocaine (bottom) injected WT and KO mice. (C2) mEPSC amplitude; (C3) mEPSC frequency; (C4-C5) mEPSC rise and decay time. Saline group: N = 13-16 neurons per genotype, 3-4 animals per genotype; cocaine group: (N = 22-27 neurons per genotype, 4-5 animals per genotype). (D1) representative traces showing mIPSCs at holding potential of +5 mV in presence of 0.5 µM TTX in saline (top) or cocaine (bottom) injected WT and KO mice. (D2) mIPSC amplitude; (D3) mIPSC frequency (D4-D5) mIPSC rise time and decay time. Saline group: N = 18-27 neurons per genotype, 3-4 animals per genotype; cocaine group: N = 25-31 neurons per genotype, 4-5 animals per genotype. Error bar represents SEM. *p<0.01.

Medium spiny neurons of accumbens shell receive inhibitory afferents from local interneurons, cortex and VTA (67, 68). There is no change in mIPSC amplitude in WT and KO with or without drug exposure (Fig 5D2). However, mIPSC frequency is higher in naïve KO neurons than naïve WT neurons, and cocaine exposure increases mIPSC frequency in KO but not WT neurons (Fig 5D3). Elevated mIPSC frequency in *Tmod2* KO could be a neurophysiological homeostatic adaptation to compensate for increased excitability of accumbens shell neurons due to modified intrinsic and excitatory synaptic properties (69). Rise and decay time of mIPSCs are not different in WT and KO and unaffected by cocaine exposure (Fig 5D4-D5). Combined, investigation of synaptic properties indicates preexisting and drug-induced differences in excitatory pre- and post-synaptic neurotransmission, along with augmented presynaptic inhibitory neurotransmission. These data provide a mechanistic underpinning for the lack of sensitization and acquisition in the IVSA paradigm.

## Discussion

Our work describes a critical role for *Tmod2* in drug-induced behavioral sensitization and voluntary drug intake. *Tmod2* has been weakly linked to risky drug use in humans and its gene expression and protein levels are dynamic upon drug exposure in animal models (35-41). We provide addiction relevant behavioral data that *Tmod2* regulates cocaine-induced locomotor sensitization and voluntary self-administration of cocaine, but not acute locomotor activation to cocaine. *Tmod2* mediates dendritic branching, spine maturation and density, and synaptic plasticity; the key components of experience-induced learning and memory including addiction (27, 28, 32). Concordantly, we find that drug-induced plasticity is affected in these knockouts with the deletion of *Tmod2* leading to an imbalance of excitatory and inhibitory signaling as well as compromised synaptic plasticity in the mesolimbic reward circuit. Combined, the behavioral and electrophysiological data indicate neuromodulations that mediate transition to addiction are compromised in the absence of *Tmod2*.

Structural and functional plasticity of synapses are the fundamental mechanisms that underlie addiction even though the role of actin in this process remains controversial (2, 8, 20). Acute or chronic cocaine exposure, as well as abstinence, facilitates dendritic spine outgrowth and spine density in reward-related brain regions such as prefrontal cortex and nucleus accumbens (23, 66, 70). Our previous study established a role of *Cyfip2*, a component of the WAVE regulatory complex that mediates branched actin formation in cocaine responses. We demonstrated that a *Cyfip2* mutation that exists in certain C57BL/6 substrains leads to altered acute and sensitized responses to cocaine (13). The role of actin-regulating *Tmod2* in addiction related behaviors has previously been unclear. Here we report that *Tmod2* KO mice exhibit endophenotypes that are predictive of addiction, but surprisingly exhibit resistance to drug-induced behaviors and associated neuroadaptations. Typically, hyperactivity, novelty-reactivity, anxiety, and risk-taking behaviors are associated with increased risk of addiction liability in humans and animal models (43, 71). However, the lack of *Tmod2* seems to be protective, potentially due to modifications in intrinsic and synaptic properties of reward regions that regulate reward learning (72, 73). In animal models, preexisting excitability or optogenetic stimulation of prefrontal cortex, and deep brain stimulation of accumbens shell but not core attenuates cocaine-induced behaviors (74-77). It is intriguing to speculate that the hyperexcitability of prefrontal cortex and ventral striatum neurons that we observed in *Tmod2* mutants may actively resist the cocaine-induced depression of these regions, thereby masking the maladaptive behavioral effects of cocaine. Basal and cocaine-induced changes in synaptic properties of accumbens neurons are also perturbed in *Tmod2* KO. Specifically, an elevated basal amplitude of excitatory glutamatergic synapses and unaffected release probability from presynaptic neurons in *Tmod2* KO striatal neurons demonstrate altered AMPA receptor number or function, and absence of cocaine-induced presynaptic plasticity (78, 79). Similarly, accumbens shell neurons exhibit modified, drug-independent inhibitory presynaptic responses that is further amplified by chronic drug exposure. Although drug-induced excitatory synaptic events are well characterized in nucleus accumbens, little is known about inhibitory synapses and how they respond to repeated cocaine challenge. Previous reports suggest an absence or decrease in the postsynaptic and presynaptic strengthening of inhibitory neurotransmission following cocaine exposure (79). Although available literature is biased towards the role of TMOD2 in the postsynaptic dendritic compartment, our study clearly shows presynaptic effects on neurotransmission that leads to frequency modulation in excitatory and inhibitory synapses. Interestingly, the acute locomotor response to cocaine is not affected in *Tmod2* KO and argues that the basal ganglia circuit, which regulates drug-induced locomotor activation, is not perturbed (80).

Recent human GWAS studies link *Tmod2* variants with intellectual disability (33) and intelligence (34), and suggest an important role of *Tmod*2 gene in influencing normal cognitive functions in human. Operant learning, as accessed by operant cocaine and food self-administration, is a form of associative learning that is dependent on various brain regions including nucleus accumbens and hippocampus (81, 82). *Tmod2* KO shows deficits in associative learning with delayed performance in operant conditioning, but no deficits in motor learning as determined by the rotarod assay. In addition, KO mice show enhanced non-associative learning which is demonstrated by enhanced habituation of hyperactivity and anxiety measures in the open field. Thus, there is specificity in the type of learning that is regulated by *Tmod2*.

Since *Tmod2* regulates actin function (31), we tested the mutants for structural anomalies of the brain using MRI. We found a ∼9% reduction of total brain volume, with 60% of the structures evaluated across the brain affected. However, after correction for differences in whole brain size (accounting for global volume differences), a smaller number of key structural differences were isolated: (1) decreased volumes across a number of regions notably including white matter structures and the thalamus; and (2) volume increases over a number of regions including the basal forebrain and the nucleus accumbens. Comparable human MRI studies of addiction include reports of a number of morphological or microstructural changes related to the regions identified in the *Tmod2* KO mice. For example, decreased fractional anisotropy, indicates abnormality in the white matter, and has been reported in the frontal white matter (83) and in the corpus callosum (84) of cocaine-dependent individuals. The thalamus, frequently linked to addiction, has also been studied frequently, but morphological findings are inconsistent, with volume increases, decreases or no change reported (85). More consistent findings have been observed in the nucleus accumbens, where decreased volumes have been identified in crack-cocaine users (85) and increased volumes are associated with the duration of abstinence (from alcohol, cannabis, and/or cocaine) in former users (86). It remains unclear in these human studies, however, whether the morphological findings predate or emerge after substance use. Given that the *Tmod2* KO mice evaluated by MRI in this study were cocaine naïve, it is difficult to relate the morphological findings directly to human observations. Although we did not perform measurements of fractional anisotropy in this work, the smaller white matter volumes in the *Tmod2* KO mice may be inconsistent with changes observed in humans, in which microstructural changes evident in fractional anisotropy were linked with drug use. On the other hand, the larger relative volume of the nucleus accumbens in *Tmod2* KO mice, and the corresponding reduced IVSA, appears consistent with the expectations from human findings that an increased accumbens volume would be associated with a reduced addiction phenotype (86). Of course, the morphological changes observed in the *Tmod2* KO mice represent the net result of a complex developmental process, including influences of altered synapse formation and/or function. Thus, like the behavioral phenotype, morphological changes may be secondary to other dysfunctional cellular processes. The complex interplay of these interactions is an ongoing topic in neurodevelopmental research.

The neuronal restriction of *Tmod2* expression combined with its functional role in regulating cocaine intake could potentially be harnessed as a pharmacotherapy for addiction-related behaviors. Recently, non-muscle myosin IIB which regulates synaptic actin polymerization has been proposed as a potential therapeutic target for treatment of methamphetamine addiction, but was found to be ineffective against cocaine or morphine associated contextual memory (21, 87). Our behavioral and electrophysiology data argues that *Tmod2* has a protective role in transition to addiction, perhaps through regulation of basal or drug induced balance of excitatory and inhibitory networks. The impairment in cocaine-based reward learning but no deficits in motor learning or non-associative learning, indicates specificity of TMOD2 function. Future studies with cell type and temporally specific *Tmod2* knockouts are required to tease apart this specificity and to address the developmental role of TMOD2 in the reward circuit. The current findings combined with our previous results (13) emphasize the advantage of unbiased forward genetic approaches to studying addiction-related behavior.

## Supporting information

Experimental procedure, supplementary figures S1-S3 and supplemental table S2

## Acknowledgements

We thank the members of the Kumar laboratory for their valuable input to this work especially Brain Geuther and Kevin Seburn for proofreading the manuscript and suggesting the layout of the figures. We also thank members of Dr. Zhong-Wei lab especially Dr. Guoqiang Hou for helping AM in setting up the electrophysiology rig for slice recordings. We thank Avery Lopez for her help with behavioral assay.. We thank Delia Hartley for her help with animal husbandry. We thank the entire KOMP team at The Jackson Laboratory – Jacqui White, Steve A. Murray, Robert E. Braun, James Clark, Pamelia Fraungruber, Rose Presby, Zachery Seavey, and Catherine Witmeyer. We also thank Heidi Munger and Jeremy Charette of Genome Technologies division at the Jackson Laboratory for their help in generating the RNA sequencing data. This work was supported by the NIDA grant the NIH National Institute on Drug Abuse 5P50DA039841 to E.J.C., U01DA041668 and Brain and Behavioral Foundation Young Investigator Award to V.K.

## References

1. Grueter BA, Rothwell PE, & Malenka RC (2012) Integrating synaptic plasticity and striatal circuit function in addiction. Current opinion in neurobiology 22(3):545–551.

2. Rothenfluh A & Cowan CW (2013) Emerging roles of actin cytoskeleton regulating enzymes in drug addiction: actin or reactin’? Current opinion in neurobiology 23(4):507–512.

3. Volkow Nora D & Morales M (2015) The Brain on Drugs: From Reward to Addiction. Cell 162(4):712–725.

4. Miller EC, et al. (2012) Differential modulation of drug-induced structural and functional plasticity of dendritic spines. Molecular pharmacology 82(2):333–343.

5. Kourrich S, Klug JR, Mayford M, & Thomas MJ (2012) AMPAR-independent effect of striatal αCaMKII promotes the sensitization of cocaine reward. The Journal of neuroscience : the official journal of the Society for Neuroscience 32(19):6578–6586.

6. Thomas MJ, Beurrier C, Bonci A, & Malenka RC (2001) Long-term depression in the nucleus accumbens: a neural correlate of behavioral sensitization to cocaine. Nature neuroscience 4(12):1217–1223.

7. Peoples LL, Kravitz AV, & Guillem K (2007) The role of accumbal hypoactivity in cocaine addiction. TheScientificWorldJournal 7:22–45.

8. Russo SJ, et al. (2010) The addicted synapse: mechanisms of synaptic and structural plasticity in nucleus accumbens. Trends in neurosciences 33(6):267–276.

9. Deller T, et al. (2003) Synaptopodin-deficient mice lack a spine apparatus and show deficits in synaptic plasticity. Proc Natl Acad Sci U S A 100(18):10494–10499.

10. Feng J, et al. (2000) Spinophilin regulates the formation and function of dendritic spines. Proc Natl Acad Sci U S A 97(16):9287–9292.

11. Cox PR & Zoghbi HY (2000) Sequencing, expression analysis, and mapping of three unique human tropomodulin genes and their mouse orthologs. Genomics 63(1):97–107.

12. Star EN, Kwiatkowski DJ, & Murthy VN (2002) Rapid turnover of actin in dendritic spines and its regulation by activity. Nature neuroscience 5(3):239–246.

13. Kumar V, et al. (2013) C57BL/6N mutation in cytoplasmic FMRP interacting protein 2 regulates cocaine response. Science (New York, N.Y.) 342(6165):1508–1512.

14. Pilo Boyl P, et al. (2007) Profilin2 contributes to synaptic vesicle exocytosis, neuronal excitability, and novelty-seeking behavior. The EMBO journal 26(12):2991–3002.

15. Li G, et al. (2015) Inhibition of actin polymerization in the NAc shell inhibits morphine-induced CPP by disrupting its reconsolidation. Sci Rep 5:16283.

16. Toda S, Shen HW, Peters J, Cagle S, & Kalivas PW (2006) Cocaine increases actin cycling: effects in the reinstatement model of drug seeking. The Journal of neuroscience : the official journal of the Society for Neuroscience 26(5):1579–1587.

17. Young EJ, Briggs SB, & Miller CA (2015) The Actin Cytoskeleton as a Therapeutic Target for the Prevention of Relapse to Methamphetamine Use. CNS & neurological disorders drug targets 14(6):731–737.

18. Ojelade SA, et al. (2015) Rsu1 regulates ethanol consumption in Drosophila and humans. Proceedings of the National Academy of Sciences 112(30):E4085.

19. Gu Z, Fonseca V, & Hai C-M (2013) Nicotinic acetylcholine receptor mediates nicotine-induced actin cytoskeletal remodeling and extracellular matrix degradation by vascular smooth muscle cells. Vascular pharmacology 58(1-2):87–97.

20. Miller EC, et al. (2012) Differential modulation of drug-induced structural and functional plasticity of dendritic spines. Molecular pharmacology 82(2):333–343.

21. Young EJ, et al. (2016) Nonmuscle myosin IIB as a therapeutic target for the prevention of relapse to methamphetamine use. Mol Psychiatry 21(5):615–623.

22. Kiraly DD, et al. (2010) Behavioral and morphological responses to cocaine require kalirin7. Biological psychiatry 68(3):249–255.

23. Norrholm SD, et al. (2003) Cocaine-induced proliferation of dendritic spines in nucleus accumbens is dependent on the activity of cyclin-dependent kinase-5. Neuroscience 116(1):19–22.

24. Pulipparacharuvil S, et al. (2008) Cocaine regulates MEF2 to control synaptic and behavioral plasticity. Neuron 59(4):621–633.

25. Weber A, Pennise CR, & Fowler VM (1999) Tropomodulin increases the critical concentration of barbed end-capped actin filaments by converting ADP.P(i)-actin to ADP-actin at all pointed filament ends. The Journal of biological chemistry 274(49):34637–34645.

26. Fischer RS & Fowler VM (2003) Tropomodulins: life at the slow end. Trends Cell Biol 13(11):593–601.

27. Omotade OF, et al. (2018) Tropomodulin Isoform-Specific Regulation of Dendrite Development and Synapse Formation. The Journal of Neuroscience.

28. Gray KT, et al. (2016) Tropomodulin isoforms utilize specific binding functions to modulate dendrite development. Cytoskeleton (Hoboken) 73(6):316–328.

29. Sussman MA, et al. (1994) Neural tropomodulin: developmental expression and effect of seizure activity. Brain research. Developmental brain research 80(1-2):45–53.

30. Watakabe A, Kobayashi R, & Helfman DM (1996) N-tropomodulin: a novel isoform of tropomodulin identified as the major binding protein to brain tropomyosin. Journal of cell science 109 (Pt 9):2299–2310.

31. Yamashiro S, Speicher KD, Speicher DW, & Fowler VM (2010) Mammalian tropomodulins nucleate actin polymerization via their actin monomer binding and filament pointed end-capping activities. The Journal of biological chemistry 285(43):33265–33280.

32. Cox PR, et al. (2003) Mice lacking tropomodulin-2 show enhanced long-term potentiation, hyperactivity, and deficits in learning and memory. Molecular and Cellular Neuroscience 23(1):1–12.

33. Davies G, et al. (2018) Study of 300,486 individuals identifies 148 independent genetic loci influencing general cognitive function. Nature communications 9(1):2098.

34. Hill WD, et al. (2019) A combined analysis of genetically correlated traits identifies 187 loci and a role for neurogenesis and myelination in intelligence. Mol Psychiatry 24(2):169–181.

35. Polimanti R, et al. (2017) S100A10 identified in a genome-wide gene x cannabis dependence interaction analysis of risky sexual behaviours. J Psychiatry Neurosci 42(4):252–261.

36. Iwazaki T, McGregor IS, & Matsumoto I (2006) Protein expression profile in the striatum of acute methamphetamine-treated rats. Brain Res 1097(1):19–25.

37. Zhu L, et al. (2016) mRNA changes in nucleus accumbens related to methamphetamine addiction in mice. Sci Rep 6:36993.

38. Lesscher HM, Houthuijzen JM, Groot Koerkamp MJ, Holstege FC, & Vanderschuren LJ (2012) Amygdala 14-3-3zeta as a novel modulator of escalating alcohol intake in mice. PloS one 7(5):e37999.

39. Oliver RJ, et al. (2018) Neuronal RNA-binding protein HuD regulates addiction-related gene expression and behavior. Genes Brain Behav 17(4):e12454.

40. Tapocik JD, et al. (2013) Neuroplasticity, axonal guidance and micro-RNA genes are associated with morphine self-administration behavior. Addict Biol 18(3):480–495.

41. Li CY, Mao X, & Wei L (2008) Genes and (common) pathways underlying drug addiction. PLoS Comput Biol 4(1):e2.

42. Ball SA, Kranzler HR, Tennen H, Poling JC, & Rounsaville BJ (1998) Personality disorder and dimension differences between type A and type B substance abusers. Journal of personality disorders 12(1):1–12.

43. Dickson PE, et al. (2015) Association of novelty-related behaviors and intravenous cocaine self-administration in Diversity Outbred mice. Psychopharmacology 232(6):1011–1024.

44. Kurbatova N, Mason JC, Morgan H, Meehan TF, & Karp NA (2015) PhenStat: A Tool Kit for Standardized Analysis of High Throughput Phenotypic Data. PloS one 10(7):e0131274.

45. Lever C, Burton S, & O’Keefe J (2006) Rearing on hind legs, environmental novelty, and the hippocampal formation. Reviews in the neurosciences 17(1-2):111–133.

46. Hascoet M, Bourin M, & Nic Dhonnchadha BA (2001) The mouse light-dark paradigm: a review. Progress in neuro-psychopharmacology & biological psychiatry 25(1):141–166.

47. Molander AC, et al. (2011) High impulsivity predicting vulnerability to cocaine addiction in rats: some relationship with novelty preference but not novelty reactivity, anxiety or stress. Psychopharmacology 215(4):721–731.

48. Robinson TE & Berridge KC (2003) Addiction. Annual review of psychology 54:25–53.

49. Vanderschuren LJ & Kalivas PW (2000) Alterations in dopaminergic and glutamatergic transmission in the induction and expression of behavioral sensitization: a critical review of preclinical studies. Psychopharmacology 151(2-3):99–120.

50. Phillips TJ (1997) Behavior genetics of drug sensitization. Critical reviews in neurobiology 11(1):21–33.

51. Steketee JD & Kalivas PW (2011) Drug wanting: behavioral sensitization and relapse to drug-seeking behavior. Pharmacological reviews 63(2):348–365.

52. Areal LB, Hamilton A, Martins-Silva C, Pires RGW, & Ferguson SSG (2019) Neuronal scaffolding protein spinophilin is integral for cocaine-induced behavioral sensitization and ERK1/2 activation. Molecular Brain 12(1):15.

53. Tolliver BK & Carney JM (1994) Sensitization to stereotypy in DBA/2J but not C57BL/6J mice with repeated cocaine. Pharmacology Biochemistry and Behavior 48(1):169–173.

54. Bolivar VJ (2009) Intrasession and intersession habituation in mice: from inbred strain variability to linkage analysis. Neurobiology of learning and memory 92(2):206–214.

55. Belin-Rauscent A, Fouyssac M, Bonci A, & Belin D (2016) How Preclinical Models Evolved to Resemble the Diagnostic Criteria of Drug Addiction. Biological psychiatry 79(1):39–46.

56. Benuck M, Lajtha A, & Reith ME (1987) Pharmacokinetics of systemically administered cocaine and locomotor stimulation in mice. The Journal of pharmacology and experimental therapeutics 243(1):144–149.

57. Bailey SA, Zidell RH, & Perry RW (2004) Relationships between organ weight and body/brain weight in the rat: what is the best analytical endpoint? Toxicologic pathology 32(4):448–466.

58. Agrawal HC, Davis JM, & Himwich WA (1968) Developmental changes in mouse brain: weight, water content and free amino acids. Journal of neurochemistry 15(9):917–923.

59. Parikshak NN, Gandal MJ, & Geschwind DH (2015) Systems biology and gene networks in neurodevelopmental and neurodegenerative disorders. Nature reviews. Genetics 16(8):441–458.

60. Srinivasan K, et al. (2016) Untangling the brain’s neuroinflammatory and neurodegenerative transcriptional responses. Nature communications 7:11295.

61. Torre D, Lachmann A, & Ma’ayan A (2018) BioJupies: Automated Generation of Interactive Notebooks for RNA-Seq Data Analysis in the Cloud. Cell systems 7(5):556–561.e553.

62. Kramer A, Green J, Pollard J, Jr., & Tugendreich S (2014) Causal analysis approaches in Ingenuity Pathway Analysis. Bioinformatics (Oxford, England) 30(4):523–530.

63. Dong Y & Nestler EJ (2014) The neural rejuvenation hypothesis of cocaine addiction. Trends in pharmacological sciences 35(8):374–383.

64. Gray KT, Kostyukova AS, & Fath T (2017) Actin regulation by tropomodulin and tropomyosin in neuronal morphogenesis and function. Mol Cell Neurosci 84:48–57.

65. Malenka RC & Nicoll RA (1999) Long-term potentiation--a decade of progress? Science (New York, N.Y.) 285(5435):1870–1874.

66. Robinson TE & Kolb B (2004) Structural plasticity associated with exposure to drugs of abuse. Neuropharmacology 47 Suppl 1:33–46.

67. Ishikawa M, et al. (2013) Exposure to cocaine regulates inhibitory synaptic transmission from the ventral tegmental area to the nucleus accumbens. The Journal of physiology 591(19):4827–4841.

68. Li Z, et al. (2018) Cell-Type-Specific Afferent Innervation of the Nucleus Accumbens Core and Shell. Frontiers in Neuroanatomy 12(84).

69. Kilman V, van Rossum MC, & Turrigiano GG (2002) Activity deprivation reduces miniature IPSC amplitude by decreasing the number of postsynaptic GABA(A) receptors clustered at neocortical synapses. The Journal of neuroscience : the official journal of the Society for Neuroscience 22(4):1328–1337.

70. Dos Santos M, et al. (2017) Rapid Synaptogenesis in the Nucleus Accumbens Is Induced by a Single Cocaine Administration and Stabilized by Mitogen-Activated Protein Kinase Interacting Kinase-1 Activity. Biological psychiatry 82(11):806–818.

71. Dellu F, Piazza PV, Mayo W, Le Moal M, & Simon H (1996) Novelty-seeking in rats--biobehavioral characteristics and possible relationship with the sensation-seeking trait in man. Neuropsychobiology 34(3):136–145.

72. Kauer JA & Malenka RC (2007) Synaptic plasticity and addiction. Nature reviews. Neuroscience 8(11):844–858.

73. Kourrich S, Calu DJ, & Bonci A (2015) Intrinsic plasticity: an emerging player in addiction. Nature reviews. Neuroscience 16(3):173–184.

74. Chang JY, Sawyer SF, Paris JM, Kirillov A, & Woodward DJ (1997) Single neuronal responses in medial prefrontal cortex during cocaine self-administration in freely moving rats. *Synapse (New York*, N.Y.) 26(1):22–35.

75. Sun W & Rebec GV (2006) Repeated Cocaine Self-Administration Alters Processing of Cocaine-Related Information in Rat Prefrontal Cortex. The Journal of Neuroscience 26(30):8004.

76. Chen BT, et al. (2013) Rescuing cocaine-induced prefrontal cortex hypoactivity prevents compulsive cocaine seeking. Nature 496(7445):359–362.

77. Vassoler FM, et al. (2013) Deep brain stimulation of the nucleus accumbens shell attenuates cocaine reinstatement through local and antidromic activation. The Journal of neuroscience : the official journal of the Society for Neuroscience 33(36):14446–14454.

78. Kourrich S, Rothwell PE, Klug JR, & Thomas MJ (2007) Cocaine Experience Controls Bidirectional Synaptic Plasticity in the Nucleus Accumbens. The Journal of Neuroscience 27(30):7921.

79. Kim J, Park B-H, Lee JH, Park SK, & Kim J-H (2011) Cell Type-Specific Alterations in the Nucleus Accumbens by Repeated Exposures to Cocaine. Biological psychiatry 69(11):1026–1034.

80. Bateup HS, et al. (2010) Distinct subclasses of medium spiny neurons differentially regulate striatal motor behaviors. Proc Natl Acad Sci U S A 107(33):14845–14850.

81. Brincat SL & Miller EK (2015) Frequency-specific hippocampal-prefrontal interactions during associative learning. Nature neuroscience 18(4):576–581.

82. Day JJ, Roitman MF, Wightman RM, & Carelli RM (2007) Associative learning mediates dynamic shifts in dopamine signaling in the nucleus accumbens. Nature neuroscience 10(8):1020–1028.

83. Lim KO, et al. (2008) Brain macrostructural and microstructural abnormalities in cocaine dependence. Drug and alcohol dependence 92(1-3):164–172.

84. Moeller FG, et al. (2005) Reduced anterior corpus callosum white matter integrity is related to increased impulsivity and reduced discriminability in cocaine-dependent subjects: diffusion tensor imaging. Neuropsychopharmacology : official publication of the American College of Neuropsychopharmacology 30(3):610–617.

85. Garza-Villarreal EA, et al. (2017) The effect of crack cocaine addiction and age on the microstructure and morphology of the human striatum and thalamus using shape analysis and fast diffusion kurtosis imaging. Translational psychiatry 7(5):e1122.

86. Korponay C, Kosson DS, Decety J, Kiehl KA, & Koenigs M (2017) Brain Volume Correlates with Duration of Abstinence from Substance Abuse in a Region-Specific and Substance-Specific Manner. Biological psychiatry. Cognitive neuroscience and neuroimaging 2(7):626–635.

87. Briggs SB, Blouin AM, Young EJ, Rumbaugh G, & Miller CA (2017) Memory disrupting effects of nonmuscle myosin II inhibition depend on the class of abused drug and brain region. Learning & memory (Cold Spring Harbor, N.Y.) 24(2):70–75.

