## Supplementary material for "*Tmod2* is a regulator of cocaine responses through control of striatal and cortical excitability, and drug-induced plasticity": Experimental procedure, supplementary figures S1-S3 and supplemental table S2

### Experimental procedures

#### Animals and drugs

Tropomodulin 2 (*Tmod2*$^{-/-}$) knockout mice, B6N(Cg)-*Tmod2*$^{tm1b(KOMP)Wtsi}$/2J (JR# 024939; referred to as *Tmod2* KO in text or KO in figures) with a LacZ reporter gene and C57BL/6NJ (JR# 005304; control referred to as WT) mice were obtained from The Jackson Laboratory, Bar Harbor. Both male and female mice were used for experiments. Mice were bred for at least two generations in an in-house colony before experimentation. Animal handling and experimental procedures were approved by Institutional Animal Care and Use Committees at The Jackson Laboratory. Cocaine hydrochloride (CAS no. 53-21-4, Sigma) was dissolved in 0.9% NaCl (vehicle) and was administered intraperitoneally.

#### Data mining from KOMP repository

Data generated by The Jackson Laboratory's KOMP phenotyping pipeline on *Tmod2* KO mice were harvested from International Mouse Phenotyping Consortium website and is publically available at http://www.mousephenotype.org/data/genes/MGI:1355335#section-associations. Details of the phenotyping pipeline, behavioral assays and parameters are available at http://www.mousephenotype.org/impress/procedures/7.

#### Locomotor response to acute injection of cocaine

All testing was carried out between Zeitgeber time 3 and 11 (ZT 3-11). Mice were habituated to handling and testing room conditions prior to behavioral assays. Mice (10 weeks old) were brought into the testing room and habituated to the open field testing room for 1 hour with white noise (75 ± 2 decibels), before introducing them to the open field. Locomotor activity was recorded using an overhead camera and Limelight software (Actimetrics, Wilmette, IL) before and after the injections. One single dose of low (10 mg/kg) or high (20 mg/kg) dose of cocaine was injected and the locomotor activity was recorded for 1 h (1).

#### Locomotor sensitization

Following 1 h habituation to testing room, mice (10 weeks old) were introduced in an open field arena and locomotor activity before and after the saline or cocaine injections were recorded. Mice received saline for 4 days and cocaine (15 mg/kg) for the next 6 days followed by 1 saline injection on day 11. A same-dose cocaine challenge on day 17 and an additional challenge on day 24 were given (see timeline in fig 2D) (1). Habituation learning is measured by comparing changes in locomotor activity between no-drug sessions (S1 and S4) as described in details previously (2).

#### Intravenous self-administration (IVSA) assay

The cocaine IVSA testing procedure and chambers have been described in detail previously (3). Mice began cocaine IVSA testing on a fixed-ratio 1 schedule at a dose of 1.0 mg/kg/infusion. Mice were tested in two-hour sessions at the same time daily seven days per week throughout the experiment. Acquisition criteria were defined as (a) ≥ 10 infusions per session on 3 consecutive or non-consecutive sessions followed by (b) 2 consecutive sessions during which number of infusions was ≥ 10, did not vary by more than 20%, and ≥ 70% of lever presses were on the active lever .

#### Sucrose Preference

Mice were singly housed for this protocol in a cage with the metal roof lid having inserts for 2-3 bottles and access to *ad libitum* chow. Baseline water consumption was assessed by providing water in 2 bottles for 4 consecutive days and measuring bottle weight in 24 h interval. At the end of 4th day, one of the bottles is filled with freshly prepared 4% sucrose solution and consumption was assessed for the next 4 days. The position of bottles were changed daily to avoid any physical preference or side bias towards the bottle. The bedding below the sipper bottles was monitored daily to account for spillage. If the bedding was found moist or wet with an unusual quantity of consumption for a day, the intake data was not included in the analysis. Preference is calculated as the percentage of the sucrose consumed over the total volume of fluid intake on day 8 (4).

#### Operant Conditioning for food reward

9-week old mice were subjected to mild food restriction to maintain 80-90% of *ad libitum* body weight during the entire 15 days of the protocol. Mice were habituated to the operant chamber (Med Associates, Inc., Fairfax, VT) for 30 min for 2-3 days with house light and fan on and levers retracted. Following habituation, levers were made accessible and the active lever press was associated with a delivery of one (FR1) 20 mg sucrose pellet (#S05550; Bio-serv, Flemington, NJ) as a reward and a time-out period of 20 s during which the house light is switched off. Inactive lever press was not associated with any reward or observable stimuli. Daily 1 h sessions were provided at the same time of the day, for 10 days. Total number of sucrose pellets earned, and the number of active and inactive lever press were recorded and statistically compared between *Tmod2* KO and WT. On the subsequent 5 days (day 11-15), an extinction paradigm was introduced where active lever press yielded no reward and the house light stays on for the entirety of the session. On day 16, a reinstatement session was provided, with conditions similar to the initial acquisition phase.

#### Magnetic resonance imaging

10-week old *Tmod2* KO (n=15) and WT (n=19) were deeply anesthetized and transcardially perfused with 30 ml, 1X PBS with 1 µl/ml heparin and 2mM ProHance (Bracco Diagnostics, Princeton, NJ) at room temperature, at a flow rate of 1 ml/min. Fixation, storage, and imaging procedures have been described in detail previously (5, 6). For this study, $T_2$-weighted RARE images were acquired on a 7-T scanner (Varian/Agilent) with imaging parameters: $TE_{eff}$=30ms, TR=350ms, 6 echoes, 4 averages, 40µm resolution, 13hour scan time. The resulting images were analyzed using an automated registration pipeline (7) in conjunction with predefined structural atlases (8). The resulting images obtained were analyzed for total brain, total gray matter and total white matter volume differences. Regional differences in structure were assessed through two different methods. In the first, we measured (absolute) volumes for each structure to evaluate the differences between genotypes, fitting with a linear model with sex, genotype and sex-genotype interaction coefficients. Significance was determined after correction for multiple comparisons through the false discovery rate (and expressed as q values) (9). In a second analysis, individual structure volumes were normalized to total brain volume so that each structure was evaluated as a percentage of total brain volume. The second approach accounts for a significant amount of the biological variability between mice (10).

#### Bulk RNA sequencing

Naïve, 10-week old *Tmod2* KO (n=3) and WT (n=3) mice were euthanized by cervical dislocation and the brains were harvested rapidly in ice-cold 1X PBS. 1 mm thick coronal sections were obtained using stainless steel brain matrix (Stoelting, Wood Dale, IL) and punches were taken from primary and secondary motor cortices, and striatum. The punchers were immediately transferred in 1.5 ml tubes containing 250 µl of RNA*later* (#AM7020, Invitrogen) and stored at 4˚C, overnight. Next day, these tubes were transferred to -80°C freezer for long-term storage. Tissues were lysed and homogenized in TRIzol Reagent (Ambion), then RNA was isolated using the miRNeasy Mini kit (Qiagen), according to manufacturers' protocols, including the optional DNase digest step. Sample concentration and quality were assessed using the Nanodrop 2000 spectrophotometer (Thermo Scientific) and the RNA 6000 Nano LabChip assay (Agilent Technologies). Libraries were prepared by the Genome Technologies core facility at The Jackson Laboratory using the KAPA mRNA HyperPrep Kit (KAPA Biosystems), according to the manufacturer's instructions. Briefly, the protocol entails isolation of polyA containing mRNA using oligo-dT magnetic beads, RNA fragmentation, first and second strand cDNA synthesis, ligation of Illumina-specific adapters containing a unique barcode sequence for each library, and PCR amplification. Libraries were checked for quality and concentration using the D5000 ScreenTape assay (Agilent Technologies) and quantitative PCR (KAPA Biosystems), according to the manufacturers' instructions. For sequencing, libraries were pooled and sequenced by the Genome Technologies core facility at The Jackson Laboratory, 75 bp single-end on the NextSeq 500 (Illumina) using NextSeq High Output Kit v2 reagents (Illumina).

Bioinformatic analysis were carried out at JAX Computational Services using established protocols (11-14). All paired-end RNASeq were subjected to *de novo* transcriptome assembly using Trinity (15-17). The *de novo* assembled transcripts will then be used as the reference to map individual reads and for the estimation of abundance using RSEM v 1.2.12 (12). IPA (Qiagen) was used for network generation across genes that are significantly differentially expressed. These objects were then overlaid onto canonical pathways developed from information contained in the IPA Knowledge Base. IPA's inbuilt network scoring algorithm will be used to rank the networks generated.

#### Electrophysiology

##### Microelectrode array (MEA) recordings on primary cortical neuron culture

Primary cortical neuron cultures were made from mice P0-P1 as described previously (18). Briefly, pups at P0-P1 (n=4, *Tmod2* KO; n=6, WT) were anesthetized on a cold metallic block placed on a bed of ice. Brains were taken out and the entire cortical region was micro-dissected in an aseptic environment (a bio-chemical hood). Cortical tissues were immediately transferred to ice cold hibernate AB complete media (BrainBits, Springfield, IL) and were shipped to collaborators at University of Illinois at Urbana-Champaign, Urbana, IL, USA. Details of cell dissociation, maintenance on MEA plates, recordings, parameters analyzed and compared was described previously (19). Briefly, on DIV 15, MEA plates were placed inside the recording chamber with inbuilt humidifier, and access to 5% $CO_2$ at 37˚C. 15 min baseline recording were obtained before application of pharmacological agents such as GABA-A receptor blocker picrotoxin (PTX) and AMPA receptor antagonist NBQX. For analysis, 10 min baseline spiking activity before drug application was compared with 10 min response window following 5 min of drug application (Fig 4B). Fold change in spontaneous spike rate, burst number and synchrony index is a ratio of response and baseline periods. Fold change of 1 represents no change, >1 represent response value of the parameter was more than baseline and <1 represents a decrease in response value.

##### Intrinsic properties of cortical and striatal neurons

Mice were deeply anesthetized using 4% tribromoethonol (800mg/kg), decapitated and the brain was quickly dissected out in ice-cold solution containing (in mM): 210 sucrose, 26 $NaHCO_3$, 10 glucose, 3 KCl, 4 $MgCl_2$, and 1 $CaCl_2$ aerated with 95% $O_2$ and 5% $CO_2$. Using a vibratome (Leica VT1200) 250 µm brain slices from naïve, 8-week old *Tmod2* KO (n=6) and WT (n=6) mice were obtained containing pre-limbic area (coronal sections) of the medial prefrontal cortex or accumbens shell (sagittal sections). Slices were then transferred to a holding chamber with artificial cerebrospinal fluid (ACSF) at room temperature containing (in mM): 124 NaCl, 3.0 KCl, 1.5 $CaCl_2$, 1.3 $MgCl_2$, 1.0 $NaH_2PO_4$, 26 $NaHCO_3$, and 20 glucose; saturated with 95% $O_2$ and 5% $CO_2$ at room temperature (21–24°C). Slices were incubated for at least 1 h for neurons to recover and equilibrate to the ACSF, before performing the whole-cell recordings. For recordings, slices were transferred to a submersion-type chamber attached to a microscope and were continuously perfused with ACSF saturated with 95% $O_2$ and 5% $CO_2$ at 30-32˚C. Pyramidal cells of the prelimbic cortex and medium spiny neurons in accumbens shell were identified using 40X objective. Medium spiny neurons were additionally screened by their low resting membrane potential of -75 mV or lower. Whole-cell recordings were obtained from neuron cell bodies using MultiClamp 700B amplifier (Molecular Devices, Sunnyvale, CA, United States) and glass pipette containing (in mM): 120 K-gluconate, 20 KCl, 10 HEPES, 0.2 EGTA, 2 $MgCl_2$, and ATP/GTP (pH 7.2-7.3). Series resistance was fully compensated in current clamp mode. All the protocols were made and executed through the AxoGraph X software (AxoGraph Scientific). Data were filtered at 4 kHz and digitized at 20 kHz. Data analysis was performed using AxoGraph X.

Intrinsic properties of neurons were analyzed using methods described previously (20, 21). Resting membrane potentials were measured within 20 s of break-in. Input resistance, time constant and whole-cell capacitance were calculated from voltage responses to 700 ms current steps of -40 pA. To compensate for changes in resting membrane potential, cells in prelimbic cortex and accumbens shell were held at -70 and -80 mV, respectively, between current steps.

#### Synaptic properties of striatal neurons

9-week old *Tmod2* KO (n=6) and WT (n=6) were either injected with saline (n=6, 3 from each strain) or cocaine (15mg/kg; n=6, 3 from each strain), once per day for 5 days. Following injections, mice were transferred to a new cage to facilitate locomotor sensitization (22). After 3-8 days of abstinence, mice were sacrificed and sagittal brain sections containing accumbens shell regions were obtained as mentioned above. The recording ACSF contains 0.5 µM tetrodotoxin in addition to the ingredients mentioned earlier. A non-potassium, cesium (Cs) based pipette solution was used containing (in mM): 117 Cs-gluconate, 20.0 HEPES, 0.4 EGTA, 2.8 NaCl, 5.0 tetraethylammonium chloride and 4.0 QX-314 (pH 7.2-7.3). Recordings were performed in voltage-clamp mode and cells were held at -80 mV for recording miniature excitatory postsynaptic current (mEPSC) and thereafter, at +5 mV for recording miniature inhibitory postsynaptic currents (mIPSC). Amplitude, frequency, rise and decay time of mEPSCs and mIPSCs were analysed using AxoGraph X as described previously (23).

#### Conflict of Interest

We have no conflicts of interest.

#### Author contributions

Conceptualization, VK and AM; Investigation, AM, SD, PED, JZ, JG, BJN, N-PT and VK; Writing, AM and VK; Editing and review, PED, JZ, BJN, RMH, N-PT, EJC and Z-WZ; Graphs and figure layout, AM, VK, JZ, N-PT and BJN; Funding acquisition, VK; Supervision, VK and Z-WZ.

#### References


1. Kumar V, *et al.* (2011) Second-generation high-throughput forward genetic screen in mice to isolate subtle behavioral mutants. *Proceedings of the National Academy of Sciences* 108(Supplement 3):15557.
2. Bolivar VJ (2009) Intrasession and intersession habituation in mice: from inbred strain variability to linkage analysis. *Neurobiology of learning and memory* 92(2):206-214.
3. Dickson PE, *et al.* (2011) Genotype-dependent effects of adolescent nicotine exposure on dopamine functional dynamics in the nucleus accumbens shell in male and female mice: a potential mechanism underlying the gateway effect of nicotine. *Psychopharmacology* 215(4):631-642.
4. Eagle AL, *et al.* (2015) Experience-Dependent Induction of Hippocampal DeltaFosB Controls Learning. *The Journal of neuroscience : the official journal of the Society for Neuroscience* 35(40):13773-13783.
5. de Guzman AE, Wong MD, Gleave JA, & Nieman BJ (2016) Variations in post-perfusion immersion fixation and storage alter MRI measurements of mouse brain morphometry. *NeuroImage* 142:687-695.
6. Spencer Noakes TL, Henkelman RM, & Nieman BJ (2017) Partitioning k-space for cylindrical three-dimensional rapid acquisition with relaxation enhancement imaging in the mouse brain. *NMR in biomedicine* 30(11).
7. Friedel M, van Eede MC, Pipitone J, Chakravarty MM, & Lerch JP (2014) Pydpiper: a flexible toolkit for constructing novel registration pipelines. *Frontiers in neuroinformatics* 8:67.
8. Dorr AE, Lerch JP, Spring S, Kabani N, & Henkelman RM (2008) High resolution three-dimensional brain atlas using an average magnetic resonance image of 40 adult C57Bl/6J mice. *NeuroImage* 42(1):60-69.
9. Benjamini Y & Hochberg Y (1995) Controlling the False Discovery Rate: A Practical and Powerful Approach to Multiple Testing. *Journal of the Royal Statistical Society. Series B (Methodological)* 57(1):289-300.
10. Lerch JP, *et al.* (2012) Wanted dead or alive? The tradeoff between in-vivo versus ex-vivo MR brain imaging in the mouse. *Frontiers in neuroinformatics* 6:6.
11. Barnett DW, Garrison EK, Quinlan AR, Stromberg MP, & Marth GT (2011) BamTools: a C++ API and toolkit for analyzing and managing BAM files. *Bioinformatics (Oxford, England)* 27(12):1691-1692.
12. Li B & Dewey CN (2011) RSEM: accurate transcript quantification from RNA-Seq data with or without a reference genome. *BMC bioinformatics* 12:323.
13. Patel RK & Jain M (2012) NGS QC Toolkit: a toolkit for quality control of next generation sequencing data. *PloS one* 7(2):e30619.
14. Robinson MD, McCarthy DJ, & Smyth GK (2010) edgeR: a Bioconductor package for differential expression analysis of digital gene expression data. *Bioinformatics (Oxford, England)* 26(1):139-140.
15. Davies G, *et al.* (2018) Study of 300,486 individuals identifies 148 independent genetic loci influencing general cognitive function. *Nature communications* 9(1):2098.
16. Grabherr MG, *et al.* (2011) Full-length transcriptome assembly from RNA-Seq data without a reference genome. *Nature biotechnology* 29(7):644-652.
17. Haas BJ, *et al.* (2013) De novo transcript sequence reconstruction from RNA-seq using the Trinity platform for reference generation and analysis. *Nature protocols* 8(8):1494-1512.

18. Tsai NP*, et al.* (2012) Multiple autism-linked genes mediate synapse elimination via proteasomal degradation of a synaptic scaffold PSD-95. *Cell* 151(7):1581-1594.
19. Jewett KA, Lee KY, Eagleman DE, Soriano S, & Tsai N-P (2018) Dysregulation and restoration of homeostatic network plasticity in fragile X syndrome mice. *Neuropharmacology* 138:182-192.
20. Zhang ZW (2004) Maturation of layer V pyramidal neurons in the rat prefrontal cortex: intrinsic properties and synaptic function. *Journal of neurophysiology* 91(3):1171-1182.
21. Zhang ZW & Arsenault D (2005) Gain modulation by serotonin in pyramidal neurones of the rat prefrontal cortex. *The Journal of physiology* 566(Pt 2):379-394.
22. Badiani A & Robinson TE (2004) Drug-induced neurobehavioral plasticity: the role of environmental context. *Behavioural pharmacology* 15(5-6):327-339.
23. Zhang W, Peterson M, Beyer B, Frankel WN, & Zhang Z-w (2014) Loss of MeCP2 From Forebrain Excitatory Neurons Leads to Cortical Hyperexcitation and Seizures. *The Journal of Neuroscience* 34(7):2754.

Supplementary figure S1

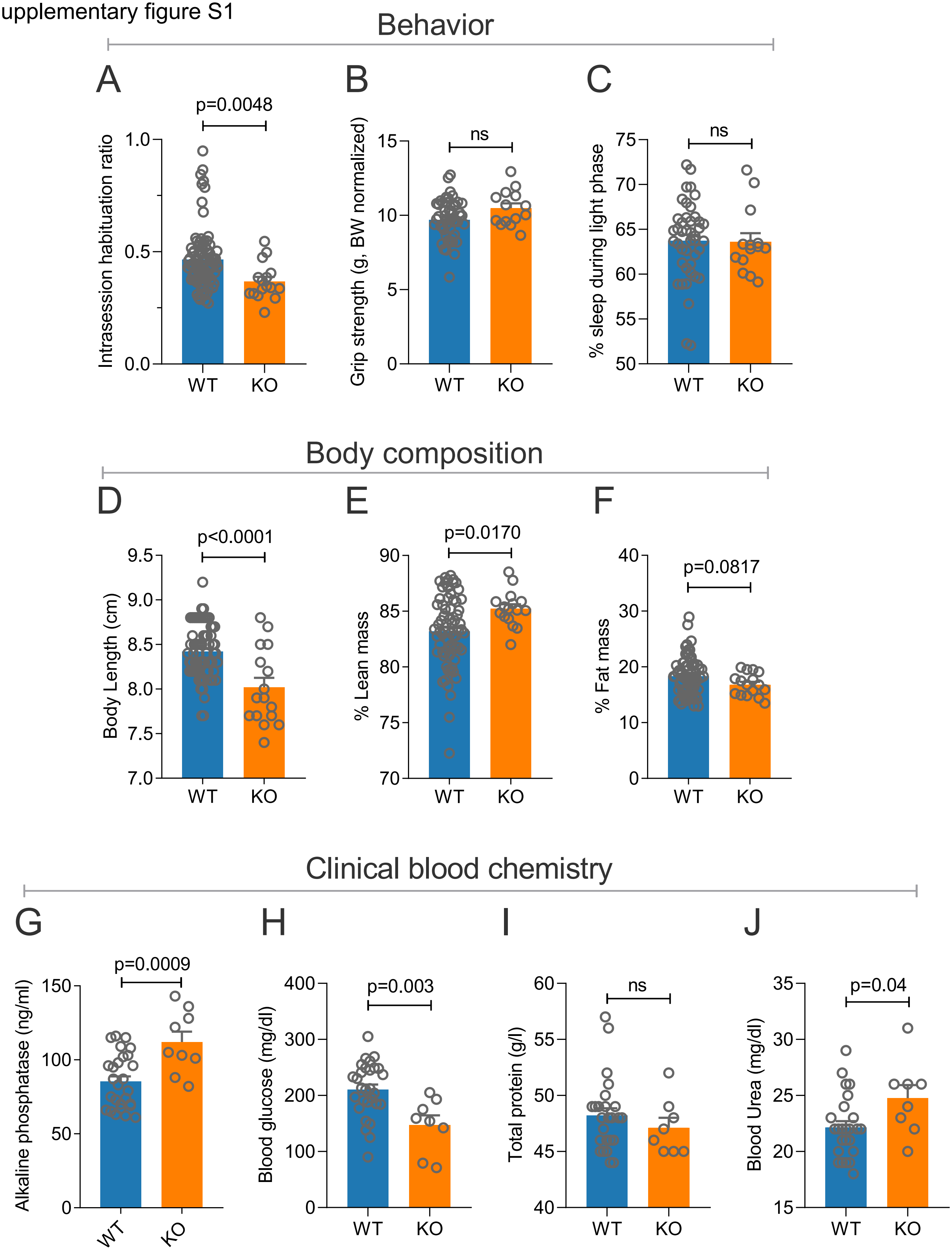

Supplementary figure S2

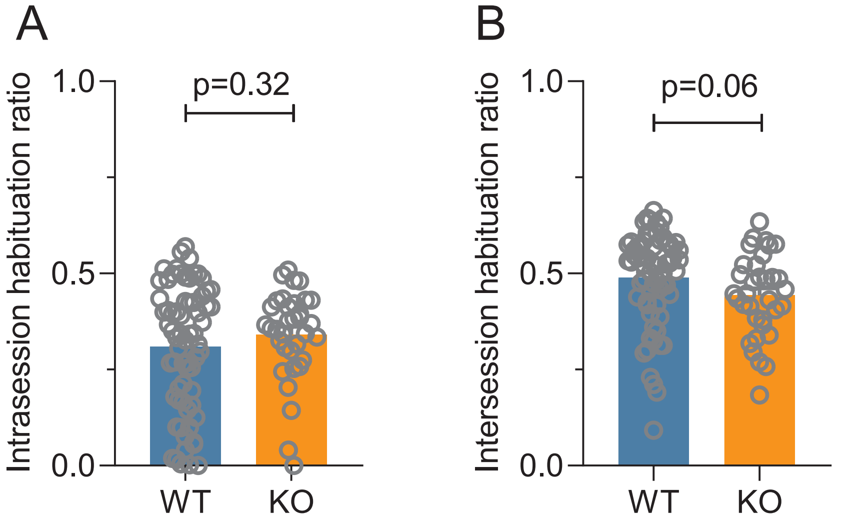

Supplementary figure S3

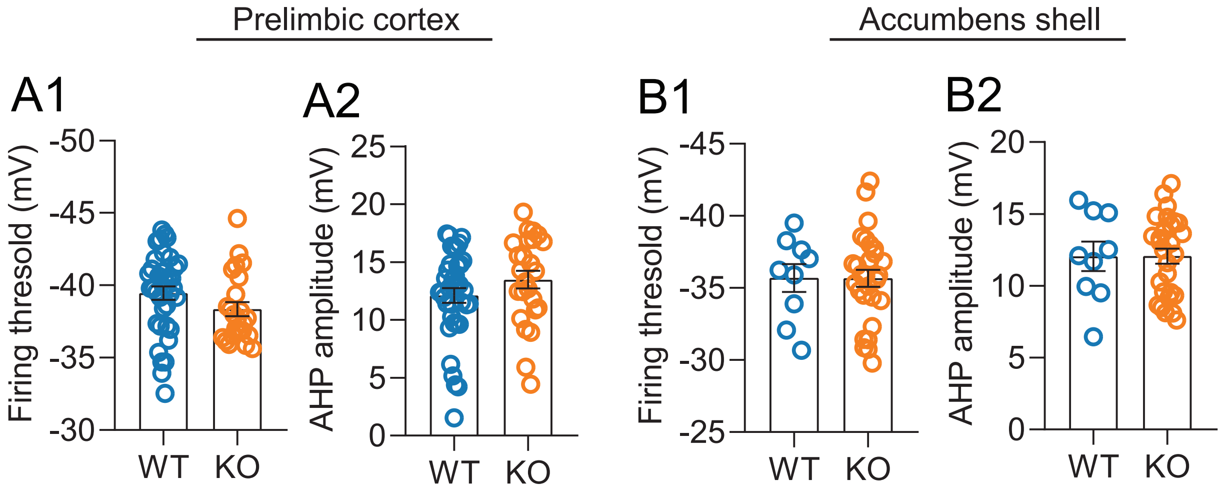

Supplementary table S2 - Summary of absoulte brain volume of different brain regions using magnetic resonance imaging.

| | V.B6NJ | dV.TMOD2 | t.TMOD2 | q.TMOD2 | dV.Male | t.Male | q.Male |
|---|---|---|---|---|---|---|---|
| amygdala | 9.28 | -0.636 | -2.17 | 0.0597 | 0.355 | 1.35 | 0.71 |
| anterior commissure: pars anterior | 0.933 | -0.108 | -4.03 | 0.00122 | 0.021 | 0.872 | 0.821 |
| anterior commissure: pars posterior | 0.394 | -0.118 | -7.01 | 1.55e-06 | 0.0141 | 0.936 | 0.811 |
| basal forebrain | 4.56 | -0.214 | -1.88 | 0.1 | 0.142 | 1.39 | 0.685 |
| bed nucleus of stria terminalis | 1.22 | 0.00591 | 0.134 | 0.904 | 0.144 | 3.66 | 0.0882 |
| cerebellar peduncle: inferior | 0.791 | -0.069 | -1.73 | 0.131 | -0.0569 | -1.58 | 0.62 |
| cerebellar peduncle: middle | 1.18 | -0.119 | -3.5 | 0.00387 | -0.0918 | -3.02 | 0.157 |
| cerebellar peduncle: superior | 0.886 | -0.0297 | -1.18 | 0.291 | 0.0229 | 1.01 | 0.771 |
| cerebral aqueduct | 0.472 | -0.0408 | -0.868 | 0.446 | -0.0322 | -0.762 | 0.821 |
| cerebral peduncle | 1.98 | -0.513 | -12.3 | 5.14e-11 | -0.0189 | -0.505 | 0.878 |
| colliculus: inferior | 4.76 | -0.214 | -2.26 | 0.0517 | 0.0561 | 0.659 | 0.844 |
| colliculus: superior | 7.34 | -0.431 | -2.23 | 0.0532 | 0.264 | 1.52 | 0.62 |
| corpus callosum | 11.7 | -3.02 | -9.17 | 2e-08 | 0.0702 | 0.238 | 0.93 |
| corticospinal tract/pyramids | 1.27 | -0.183 | -2.49 | 0.0327 | -0.0682 | -1.04 | 0.766 |
| cuneate nucleus | 0.226 | -0.0282 | -1.44 | 0.203 | -0.00755 | -0.429 | 0.892 |
| facial nerve (cranial nerve 7) | 0.204 | -0.0112 | -0.758 | 0.511 | -0.0113 | -0.853 | 0.821 |
| fasciculus retro flexus | 0.221 | -0.017 | -3.48 | 0.00394 | 0.000394 | 0.09 | 0.977 |
| fimbria | 3.01 | -0.446 | -3.56 | 0.00343 | 0.0769 | 0.685 | 0.825 |
| fornix | 0.583 | -0.0885 | -6.04 | 1.51e-05 | 0.00988 | 0.751 | 0.821 |
| fourth ventricle | 0.832 | -0.0268 | -0.482 | 0.674 | -0.0602 | -1.21 | 0.71 |
| fundus of striatum | 0.157 | -0.0198 | -2.3 | 0.0477 | 0.00614 | 0.794 | 0.821 |
| globus pallidus | 3.32 | -0.554 | -3.36 | 0.00512 | 0.0747 | 0.505 | 0.878 |
| habenular commissure | 0.0299 | -0.00228 | -0.68 | 0.55 | -0.00453 | -1.5 | 0.62 |
| hypothalamus | 9.61 | -0.421 | -2.23 | 0.0532 | 0.18 | 1.06 | 0.761 |
| inferior olivary complex | 0.316 | -0.0507 | -2.15 | 0.0612 | -0.025 | -1.18 | 0.71 |
| internal capsule | 3.1 | -0.609 | -7.5 | 4.74e-07 | 0.0371 | 0.509 | 0.878 |
| interpeduncular nucleus | 0.25 | -0.00616 | -0.63 | 0.582 | -0.00565 | -0.643 | 0.845 |
| lateral olfactory tract | 1.38 | -0.0438 | -1.64 | 0.149 | 0.0372 | 1.55 | 0.62 |
| lateral septum | 2.9 | -0.0957 | -0.944 | 0.404 | 0.0405 | 0.445 | 0.892 |
| lateral ventricle | 3.7 | -0.28 | -1.06 | 0.352 | -0.104 | -0.437 | 0.892 |
| mammillary bodies | 0.424 | -0.0424 | -2.95 | 0.0125 | -0.00438 | -0.339 | 0.93 |
| mammilothalamic tract | 0.221 | -0.0354 | -5.71 | 2.35e-05 | -0.00137 | -0.247 | 0.93 |
| [illegible]edial lemniscus/medial longitudinal fasciculus | 2.08 | -0.183 | -1.88 | 0.1 | -0.0633 | -0.723 | 0.821 |
| medial septum | 1.07 | -0.054 | -1.95 | 0.0883 | 0.0154 | 0.618 | 0.849 |
| medulla | 23.6 | -1.73 | -1.84 | 0.108 | -0.48 | -0.569 | 0.856 |
| midbrain | 12.1 | -1.22 | -4.32 | 0.000618 | 0.114 | 0.451 | 0.892 |
| nucleus accumbens | 3.82 | -0.102 | -0.701 | 0.539 | 0.101 | 0.779 | 0.821 |
| olfactory peduncle | 2.37 | -0.00664 | -0.122 | 0.909 | 0.0451 | 0.922 | 0.814 |
| olfactory tubercle | 3.08 | -0.197 | -1.4 | 0.212 | -0.0207 | -0.164 | 0.943 |
| optic tract | 1.34 | -0.255 | -6.4 | 6.87e-06 | 0.0216 | 0.605 | 0.849 |
| periaqueductal grey | 3.57 | -0.0439 | -0.452 | 0.689 | 0.102 | 1.17 | 0.71 |
| pons | 15.2 | -0.475 | -1.29 | 0.251 | -0.0778 | -0.235 | 0.93 |
| pontine nucleus | 0.724 | -0.0799 | -2 | 0.0808 | -0.0799 | -2.22 | 0.41 |
| posterior commissure | 0.12 | -0.00375 | -1.22 | 0.278 | 0.000188 | 0.0679 | 0.984 |
| pre-para subiculum | 2.28 | -0.404 | -7.71 | 3.49e-07 | -0.0423 | -0.899 | 0.814 |
| stria medullaris | 0.623 | -0.0651 | -4.12 | 0.000999 | 0.00812 | 0.573 | 0.856 |

| | V.B6NJ | dV.TMOD2 | t.TMOD2 | q.TMOD2 | dV.Male | t.Male | q.Male |
|---|---|---|---|---|---|---|---|
| *stria terminalis* | 0.858 | −0.17 | −8.12 | 1.41e-07 | 0.0198 | 1.05 | 0.765 |
| *striatum* | 19.4 | −2.64 | −6.01 | 1.53e-05 | −0.645 | −1.63 | 0.62 |
| *subependymale zone / rhinocele* | 0.043 | 0.0031 | 2.23 | 0.0534 | 0.00144 | 1.15 | 0.726 |
| *superior olivary complex* | 0.671 | −0.0222 | −0.727 | 0.528 | −0.00762 | −0.278 | 0.93 |
| *thalamus* | 16.5 | −2.59 | −6.5 | 5.74e-06 | 0.0162 | 0.0451 | 0.992 |
| *third ventricle* | 1.15 | −0.0594 | −1.27 | 0.259 | −0.0334 | −0.794 | 0.821 |
| *ventral tegmental decussation* | 0.116 | −0.00633 | −1.46 | 0.197 | 0.00355 | 0.911 | 0.814 |
| *bules 1-2: lingula and central lobule (ventral)* | 1.76 | −0.277 | −3.97 | 0.0014 | −0.0556 | −0.889 | 0.816 |
| *lobule 3: central lobule (dorsal)* | 1.97 | −0.173 | −2.3 | 0.0477 | 0.017 | 0.252 | 0.93 |
| *lobules 4-5: culmen (ventral and dorsal)* | 4.27 | −0.348 | −3.4 | 0.00483 | −0.0893 | −0.972 | 0.791 |
| *lobule 6: declive* | 2.64 | −0.296 | −2.11 | 0.0657 | −0.152 | −1.21 | 0.71 |
| *lobule 7: tuber (or folium)* | 0.995 | −0.126 | −1.66 | 0.143 | −0.0755 | −1.11 | 0.746 |
| *lobule 8: pyramis* | 1.55 | −0.23 | −2.72 | 0.0203 | −0.0983 | −1.3 | 0.71 |
| *lobule 9: uvula* | 2.8 | −0.27 | −2 | 0.0808 | −0.0269 | −0.221 | 0.93 |
| *lobule 10: nodulus* | 1.26 | −0.116 | −1.48 | 0.192 | −0.0456 | −0.646 | 0.845 |
| *anterior lobule (lobules 4-5)* | 1.65 | −0.213 | −5.15 | 8.86e-05 | −0.0488 | −1.31 | 0.71 |
| *simple lobule (lobule 6)* | 4.75 | −0.38 | −3.64 | 0.00296 | −0.259 | −2.76 | 0.197 |
| *crus 1: ansiform lobule (lobule 6)* | 4.44 | −0.275 | −2.77 | 0.0183 | −0.235 | −2.64 | 0.238 |
| *crus 2: ansiform lobule (lobule 7)* | 4.3 | −0.411 | −3.04 | 0.0104 | −0.347 | −2.86 | 0.173 |
| *paramedian lobule (lobule 7)* | 4.09 | −0.618 | −4.18 | 0.000862 | −0.231 | −1.74 | 0.62 |
| *copula: pyramis (lobule 8)* | 2.32 | −0.183 | −1.71 | 0.133 | −0.146 | −1.52 | 0.62 |
| *flocculus (FL)* | 0.972 | −0.121 | −4.39 | 0.000532 | 0.012 | 0.485 | 0.891 |
| *paraflocculus (PFL)* | 3.63 | −0.258 | −1.43 | 0.203 | −0.0407 | −0.252 | 0.93 |
| *trunk of arbor vita* | 3.71 | −0.435 | −6.31 | 7.71e-06 | −0.00652 | −0.105 | 0.97 |
| *lobule 1-2 white matter* | 0.0637 | −0.0104 | −2.33 | 0.0459 | −0.00278 | −0.696 | 0.821 |
| *lobule 3 white matter* | 0.151 | −0.01 | −0.958 | 0.398 | −0.00201 | −0.213 | 0.93 |
| *trunk of lobules 1-3 white matter* | 0.132 | −0.0257 | −5.13 | 8.86e-05 | −0.00557 | −1.24 | 0.71 |
| *lobules 4-5 white matter* | 0.488 | −0.0059 | −0.199 | 0.862 | −0.00453 | −0.171 | 0.943 |
| *lobules 6-7 white matter* | 0.578 | −0.0391 | −1.41 | 0.21 | 0.000103 | 0.00413 | 0.997 |
| *lobule 8 white matter* | 0.148 | −0.0249 | −2.84 | 0.0155 | −0.0164 | −2.09 | 0.513 |
| *trunk of lobules 6-8 white matter* | 0.109 | −0.0142 | −3.28 | 0.00612 | 0.000836 | 0.215 | 0.93 |
| *lobule 9 white matter* | 0.252 | −0.0216 | −1.4 | 0.211 | −0.0204 | −1.48 | 0.62 |
| *lobule 10 white matter* | 0.0715 | −0.00728 | −1.61 | 0.155 | −0.00331 | −0.815 | 0.821 |
| *anterior lobule white matter* | 0.0849 | −0.0138 | −5.45 | 4.45e-05 | −0.00348 | −1.53 | 0.62 |
| *simple lobule white matter* | 0.349 | −0.0452 | −3.65 | 0.00289 | −0.0228 | −2.05 | 0.513 |
| *crus 1 white matter* | 0.359 | −0.0278 | −2.07 | 0.0705 | −0.0201 | −1.68 | 0.62 |
| *trunk of simple and crus 1 white matter* | 0.128 | −0.00647 | −1.5 | 0.188 | −0.0012 | −0.309 | 0.93 |
| *crus 2 white matter* | 0.251 | −0.0262 | −2.21 | 0.0544 | −0.0151 | −1.42 | 0.667 |
| *paramedian lobule* | 0.133 | −0.023 | −3.27 | 0.00612 | −0.0103 | −1.64 | 0.62 |
| *trunk of crus 2 and paramedian white matter* | 0.296 | −0.0284 | −3.84 | 0.00179 | −0.0106 | −1.6 | 0.62 |
| *copula white matter* | 0.0626 | −0.00779 | −2.27 | 0.0507 | −0.00502 | −1.63 | 0.62 |
| *paraflocculus white matter* | 0.272 | −0.0426 | −3.9 | 0.00162 | −0.023 | −2.35 | 0.39 |
| *flocculus white matter* | 0.065 | −0.0109 | −4.43 | 0.000495 | −0.0026 | −1.18 | 0.71 |
| *dentate nucleus* | 0.357 | −0.0445 | −4.78 | 0.000201 | 0.0037 | 0.443 | 0.892 |
| *nucleus interpositus* | 0.419 | −0.0519 | −4.05 | 0.00115 | 0.0222 | 1.94 | 0.568 |
| *fastigial nucleus* | 0.449 | −0.0539 | −3.87 | 0.0017 | 0.0237 | 1.9 | 0.585 |

| | V.B6NJ | dV.TMOD2 | t.TMOD2 | q.TMOD2 | dV.Male | t.Male | q.Male |
|---|---|---|---|---|---|---|---|
| *Cingulate cortex: area 24a* | 2.07 | −0.187 | −1.49 | 0.19 | 0.0456 | 0.403 | 0.897 |
| *Cingulate cortex: area 24a'* | 0.936 | −0.137 | −4.45 | 0.000479 | −0.0166 | −0.602 | 0.849 |
| *Cingulate cortex: area 24b* | 1.54 | −0.207 | −3.58 | 0.00338 | −0.119 | −2.28 | 0.392 |
| *Cingulate cortex: area 24b'* | 0.69 | −0.133 | −5.75 | 2.35e-05 | −0.0423 | −2.04 | 0.513 |
| *Cingulate cortex: area 25* | 0.624 | −0.00643 | −0.189 | 0.866 | 0.0111 | 0.364 | 0.927 |
| *Cingulate cortex: area 29a* | 0.636 | −0.0346 | −1.23 | 0.275 | 0.00475 | 0.188 | 0.94 |
| *Cingulate cortex: area 29b* | 0.394 | −0.0282 | −1.82 | 0.11 | 0.0101 | 0.724 | 0.821 |
| *Cingulate cortex: area 29c* | 2.04 | −0.34 | −5.71 | 2.35e-05 | −0.0946 | −1.77 | 0.62 |
| *Cingulate cortex: area 30* | 2.8 | −0.467 | −4.7 | 0.000249 | −0.0383 | −0.429 | 0.892 |
| *Cingulate cortex: area 32* | 2.35 | −0.146 | −1.05 | 0.354 | 0.0242 | 0.194 | 0.94 |
| *Amygdalopiriform transition area* | 1.13 | −0.0705 | −1.72 | 0.133 | −0.0113 | −0.305 | 0.93 |
| *Primary auditory cortex* | 1.54 | −0.212 | −4.45 | 0.000479 | −0.0698 | −1.63 | 0.62 |
| *Secondary auditory cortex: dorsal area* | 1.53 | −0.165 | −3.49 | 0.00392 | −0.0385 | −0.905 | 0.814 |
| *Secondary auditory cortex: ventral area* | 1.6 | −0.263 | −3.51 | 0.00387 | −0.0478 | −0.708 | 0.821 |
| *Caudomedial entorhinal cortex* | 5.92 | −0.706 | −5.73 | 2.35e-05 | −0.124 | −1.12 | 0.746 |
| *Cingulum* | 0.84 | −0.218 | −11.9 | 6.24e-11 | 0.00572 | 0.348 | 0.93 |
| *Claustrum* | 0.279 | −0.0206 | −2.99 | 0.0114 | 0.00165 | 0.268 | 0.93 |
| *Cortex-amygdala transition zones* | 0.72 | −0.0944 | −2.53 | 0.0309 | 0.0511 | 1.52 | 0.62 |
| *Claustrum: dorsal part* | 0.295 | −0.0357 | −2.87 | 0.0146 | −0.0224 | −2.01 | 0.514 |
| *Dorsal nucleus of the endopiriform* | 1.49 | −0.139 | −2.12 | 0.065 | 0.07 | 1.19 | 0.71 |
| *Dorsal intermediate entorhinal cortex* | 1.8 | −0.151 | −3.02 | 0.0108 | −0.055 | −1.23 | 0.71 |
| *Dorsolateral entorhinal cortex* | 2.57 | −0.442 | −5.14 | 8.86e-05 | −0.0195 | −0.253 | 0.93 |
| *Dorsolateral orbital cortex* | 0.901 | −0.0998 | −2.72 | 0.0203 | −0.00172 | −0.0522 | 0.991 |
| *Dorsal tenia tecta* | 1.09 | −0.00249 | −0.0556 | 0.956 | 0.0185 | 0.459 | 0.892 |
| *Ectorhinal cortex* | 2.89 | −0.403 | −2.61 | 0.0263 | 0.000516 | 0.00371 | 0.997 |
| *Frontal cortex: area 3* | 0.837 | −0.0939 | −3.96 | 0.00143 | −0.00367 | −0.172 | 0.943 |
| *Frontal association cortex* | 6.74 | −0.648 | −3.23 | 0.00669 | −0.29 | −1.61 | 0.62 |
| *ntermediate nucleus of the endopiriform claustrum* | 0.543 | −0.0642 | −2.5 | 0.0327 | 0.0227 | 0.984 | 0.787 |
| *Insular region: not subdivided* | 8.23 | −1.04 | −3.73 | 0.00239 | −0.197 | −0.786 | 0.821 |
| *Lateral orbital cortex* | 3.55 | −0.477 | −4.31 | 0.000626 | −0.0616 | −0.62 | 0.849 |
| *Lateral parietal association cortex* | 0.25 | −0.0357 | −3.19 | 0.00729 | −0.00604 | −0.601 | 0.849 |
| *Primary motor cortex* | 7.48 | −0.872 | −4.79 | 0.000201 | −0.267 | −1.63 | 0.62 |
| *Secondary motor cortex* | 6.36 | −0.64 | −3.43 | 0.00449 | −0.209 | −1.25 | 0.71 |
| *Medial entorhinal cortex* | 0.789 | −0.0414 | −2.04 | 0.075 | −0.0183 | −1.01 | 0.771 |
| *Medial orbital cortex* | 1.92 | −0.166 | −1.66 | 0.143 | −0.00681 | −0.0759 | 0.983 |
| *Medial parietal association cortex* | 0.406 | −0.072 | −3.56 | 0.00343 | −0.00462 | −0.254 | 0.93 |
| *Piriform cortex* | 10.3 | −1.14 | −2.53 | 0.0309 | 0.229 | 0.568 | 0.856 |
| *Posterolateral cortical amygdaloid area* | 0.866 | −0.0512 | −1.39 | 0.214 | 0.107 | 3.22 | 0.139 |
| *Posteromedial cortical amygdaloid area* | 1.13 | −0.0659 | −1.68 | 0.14 | 0.028 | 0.797 | 0.821 |
| *Perirhinal cortex* | 2.66 | −0.367 | −2.98 | 0.0115 | 0.0455 | 0.412 | 0.897 |
| *Parietal cortex: posterior area: rostral part* | 0.121 | −0.0185 | −3.86 | 0.00171 | −0.00315 | −0.732 | 0.821 |
| *Rostral amygdalopiriform area* | 0.451 | −0.0657 | −2.99 | 0.0115 | 0.0209 | 1.06 | 0.761 |
| *Primary somatosensory cortex* | 4.33 | −0.564 | −4.8 | 0.000201 | −0.0492 | −0.466 | 0.892 |
| *Primary somatosensory cortex: barrel field* | 10.2 | −1.57 | −5.86 | 1.88e-05 | −0.808 | −3.37 | 0.128 |
| *Primary somatosensory cortex: dysgranular zone* | 0.434 | −0.0787 | −7.53 | 4.74e-07 | −0.0269 | −2.86 | 0.173 |
| *Primary somatosensory cortex: forelimb region* | 4.33 | −0.729 | −5.99 | 1.53e-05 | −0.335 | −3.07 | 0.157 |

| | V.B6NJ | dV.TMOD2 | t.TMOD2 | q.TMOD2 | dV.Male | t.Male | q.Male |
|---|---|---|---|---|---|---|---|
| *Primary somatosensory cortex: hindlimb region* | 2.51 | −0.368 | −5.12 | 8.86e-05 | −0.108 | −1.68 | 0.62 |
| *Primary somatosensory cortex: jaw region* | 0.677 | −0.109 | −5.11 | 8.98e-05 | −0.0194 | −1.01 | 0.771 |
| *Primary somatosensory cortex: shoulder region* | 0.269 | −0.0526 | −5.54 | 3.68e-05 | −0.021 | −2.46 | 0.328 |
| *Primary somatosensory cortex: trunk region* | 0.549 | −0.0885 | −4.3 | 0.000626 | −0.0133 | −0.722 | 0.821 |
| *Primary somatosensory cortex: upper lip region* | 5.98 | −0.955 | −6.38 | 6.87e-06 | −0.168 | −1.25 | 0.71 |
| *Secondary somatosensory cortex* | 6.63 | −1.08 | −3.92 | 0.00156 | −0.379 | −1.53 | 0.62 |
| *Temporal association area* | 3.29 | −0.494 | −2.9 | 0.0138 | −0.0912 | −0.596 | 0.849 |
| *Primary visual cortex* | 2.48 | −0.436 | −4.32 | 0.000618 | −0.137 | −1.51 | 0.62 |
| *Primary visual cortex: binocular area* | 2.06 | −0.349 | −5.45 | 4.45e-05 | −0.0855 | −1.48 | 0.62 |
| *Primary visual cortex: monocular area* | 1.79 | −0.248 | −3.29 | 0.00606 | −0.0182 | −0.269 | 0.93 |
| *Secondary visual cortex: lateral area* | 2.94 | −0.418 | −4.81 | 0.000199 | −0.0849 | −1.09 | 0.753 |
| *Secondary visual cortex: mediolateral area* | 0.975 | −0.118 | −2.25 | 0.0518 | −0.00139 | −0.0294 | 0.993 |
| *Secondary visual cortex: mediomedial area* | 1.66 | −0.241 | −3.13 | 0.00833 | −0.0383 | −0.555 | 0.863 |
| *Claustrum: ventral part* | 0.618 | −0.0461 | −2.4 | 0.0397 | −0.0145 | −0.839 | 0.821 |
| *Ventral nucleus of the endopiriform claustrum* | 0.508 | −0.0612 | −3.33 | 0.0056 | −0.00855 | −0.518 | 0.878 |
| *Ventral intermediate entorhinal cortex* | 1.19 | −0.0769 | −2.34 | 0.0445 | −0.036 | −1.22 | 0.71 |
| *Ventral orbital cortex* | 1.62 | −0.231 | −3.6 | 0.00326 | −0.0155 | −0.269 | 0.93 |
| *Ventral tenia tecta* | 0.118 | −0.00412 | −0.707 | 0.538 | 0.00437 | 0.834 | 0.821 |
| *CA1Or* | 2.17 | −0.496 | −8.26 | 1.18e-07 | 0.0287 | 0.533 | 0.878 |
| *LMol* | 2.3 | −0.271 | −5.86 | 1.88e-05 | 0.0515 | 1.24 | 0.71 |
| *CA1Rad* | 2.63 | −0.3 | −5.37 | 5.12e-05 | 0.114 | 2.28 | 0.392 |
| *CA2Py* | 0.195 | −0.0105 | −1.63 | 0.15 | 0.00745 | 1.29 | 0.71 |
| *CA2Or* | 0.495 | −0.0859 | −5.92 | 1.77e-05 | 0.00429 | 0.33 | 0.93 |
| *CA2Rad* | 0.444 | −0.00613 | −0.304 | 0.789 | 0.0216 | 1.19 | 0.71 |
| *CA3Py Inner* | 0.14 | −0.00232 | −0.503 | 0.663 | 0.00168 | 0.405 | 0.897 |
| *CA3Py Outer* | 0.996 | −0.0833 | −3.29 | 0.00599 | 0.0159 | 0.7 | 0.821 |
| *CA3Or* | 2.78 | −0.357 | −3.39 | 0.00486 | −0.0301 | −0.318 | 0.93 |
| *CA3Rad* | 1.88 | −0.126 | −2.72 | 0.0203 | 0.0753 | 1.81 | 0.62 |
| *SLu* | 0.743 | −0.06 | −3.27 | 0.00612 | 0.0228 | 1.38 | 0.685 |
| *MoDG* | 4.02 | −0.494 | −5.24 | 7.21e-05 | 0.0627 | 0.741 | 0.821 |
| *GrDG* | 1.19 | −0.139 | −4.07 | 0.00112 | −0.00679 | −0.221 | 0.93 |
| *PoDG* | 0.566 | −0.0464 | −2.43 | 0.0369 | −0.012 | −0.702 | 0.821 |
| *CA1Py* | 1.19 | −0.196 | −5.4 | 4.92e-05 | 0.0356 | 1.09 | 0.753 |
| *Olfactory bulb: glomerular layer* | 4.31 | −0.239 | −2.43 | 0.0369 | −0.0653 | −0.741 | 0.821 |
| *Olfactory bulb: external plexiform layer* | 6.35 | −0.164 | −1.03 | 0.359 | 0.0175 | 0.123 | 0.961 |
| *Olfactory bulb: mitral cell layer* | 1.27 | −0.0192 | −0.533 | 0.648 | 0.00108 | 0.0335 | 0.993 |
| *Olfactory bulb: internal plexiform layer* | 1.09 | −0.00839 | −0.267 | 0.813 | 0.00377 | 0.134 | 0.958 |
| *Olfactory bulb: granule cell layer* | 4.78 | 0.0581 | 0.452 | 0.689 | 0.109 | 0.943 | 0.811 |
| *cessory olfactory bulb: glomerular, external plexiform and mitral cell layer* | 0.497 | −0.0285 | −1.84 | 0.107 | −0.00196 | −0.142 | 0.957 |
| *Accessory olfactory bulb: granule cell layer* | 0.262 | 0.00439 | 0.506 | 0.663 | 0.00588 | 0.754 | 0.821 |
| *Anterior olfactory nucleus* | 2.17 | 0.055 | 0.852 | 0.453 | 0.016 | 0.276 | 0.93 |
| *subiculum* | 3.44 | −0.606 | −8.76 | 4.17e-08 | 0.000824 | 0.0133 | 0.997 |
| *Medial amygdala* | 0.957 | −0.0107 | −0.423 | 0.706 | 0.0896 | 3.94 | 0.0817 |
| *Medial preoptic nucleus* | 0.218 | 0.0039 | 0.406 | 0.715 | 0.00631 | 0.73 | 0.821 |
